# A brand-new clustering method and analysis system designed for revealing the truth of the high-dimension large data deciphered the complex composition structure of human brain endothelial cells from single-cell RNA sequence data

**DOI:** 10.1101/2023.06.14.544789

**Authors:** Boyong Wei

## Abstract

The clustering method is the key to high dimensional large data analysis, especially for single-cell NGS data in biological science and biomedicine sectors. Those data require a hierarchical clustering method to unveil important biological features including differentiation patterns, stem cell identifications, cell sub-type discovery, and so on. Traditional hierarchical clustering has several issues to be applied to large high-dimension data. There are a few new approaches invented recently trying to fill in the blank. However, these approaches were either based on low-dimension or down-sampled data after dimension reduction (Anibal et al., 2022) from methods like PCA or consumed an enormous amount of computing resources to get a massive number of layer levels with highly limited interpretable information. In order to create a practically available solution, I invented an entirely new hierarchical clustering method called the BW method which can be directly applied to high-dimension large data without a requirement for dimension reduction or massive computing resources. I applied BW clustering to six single-cell RNA sequence sample data. BW clustering brought deep insight into these sample data including sub-type, differentiation branch, cell state changes (development, aging process), and gene expression instability. BW-generated layers were very concise. For almost nineteen thousand cells, BW clustering only yielded 9 layers. An analysis system was created based on the BW clustering method which can unprecedentedly display the true form of high dimensional data space. The resource BW required is also very low as all the work done in this paper used a 16GB memory laptop only, making it easily accessible to researchers with limited computing resources. Overall, the BW clustering method represents a major advancement in high-dimensional large data analysis for biological and biomedical applications.

## Introduction

### Background

With the rising of next-generation sequence (NGS) technology, larger high-dimension data analysis methods were an urgent requirement. Nowadays, the standard procedure for single-cell NGS data is PCA (principal component analysis) based analysis pipeline as shown in the workflow chart1.

**Workflow chart 1:**
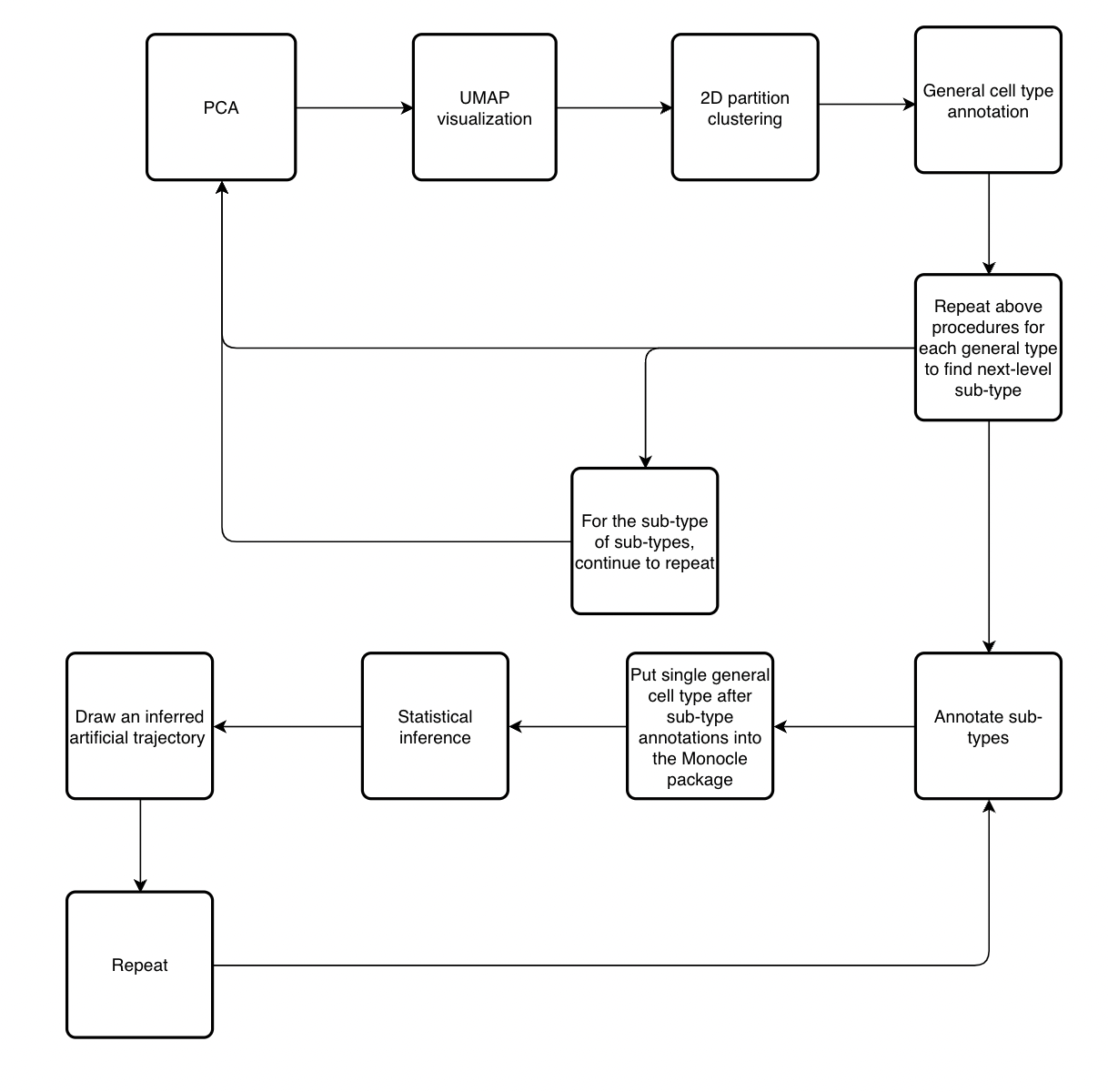
Standard single-cell analysis pipeline.

The standard procedure requires a PCA-based dimension reduction 2D graph for any further analysis. In other words, clustering in the standard procedure is made after the visualization is finished and entirely based on the visualization graph other than the data itself. On the other hand, clustering is the first step in BW based pipeline (workflow chart 2). The visualization of the BW pipeline is created on clusters generated from the clustering step. The BW pipeline only has two steps. Compared with the standard pipeline, it is a big advancement for reducing procedures.

**Workflow chart 2:**
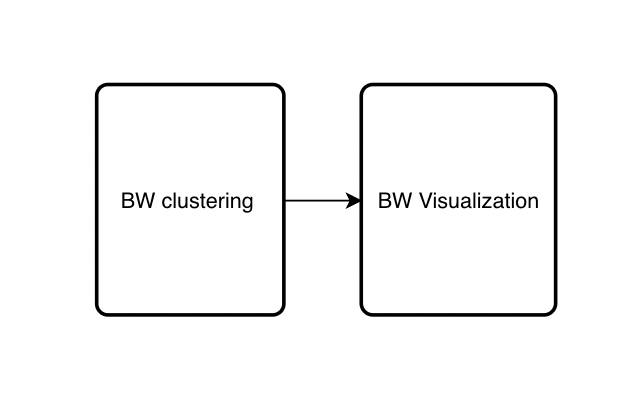
BW method pipeline.

Recently some researchers tried some other approaches to explore the hierarchical structure of single-cell data. Some of them tried to use the traditional hierarchical algorithm to deploy on single-cell NGS data (Wang et al., 2022). The main limitation of the traditional hierarchical algorithm contains two parts, and they were clearly shown in the paper (Wang et al., 2022). The first thing was extremely high-time complexity (Petegrosso et al., 2020) which means that the hierarchical tree would be unacceptably high for large data and consume an immeasurable amount of computing resources. With around 1000 cells, the number of levels could easily break through 25 levels (Wang et al., 2022). Based on the first trait of traditional hierarchical clustering, massive layers generated made interpretation extremely difficult. It is very hard to find out useful information within layer mass.

Other attempts were also reported. Some researchers tried to combine PCA and the traditional hierarchical clustering method together (žurauskiene & Yau, 2016). The pcaReduce method they made still had the same limitations. Their sample was also very small (300 or 622 cells) (žurauskiene & Yau, 2016). What is more, they tried to solve issues by setting a threshold of layers. It would be hard to say whether this kind of strategy was reliable or not. For the remaining attempts, most of them were still based on PCA-generated 2D low-dimension graphs using K-Nearest Neighbors Algorithm (KNN) (Bonald et al., 2018; Campbell et al., 2022; Seth et al., 2022) which could be entirely deviated from the reality when the data size is not small.

### The features of the BW system

The BW clustering solved all those issues, and a new analysis system was built on it. This innovative analysis procedure was tested on six samples (3 control samples, 3 disease group samples) from research on Alzheimer’s disease (Yang et al., 2022). Results showed the reality of high-dimension single-cell data space. The hierarchical structure generated from BW is not determined by sample size at all but only by the shape of high-dimensional large data. Small-size samples can run longer time than the larger sample if the form of the smaller sample is more complex. Two types of graphs were made within BW based approach: cluster graph, and trajectory graph. The cluster graph was an improved version of PCA generated UMAP reduction graph as it was created on the linear parameter. The trajectory graph created here had nothing to do with so-called trajectory graphs made by packages like Monocle in the (workflow chart 1) (Qiu et al., 2017; Van den Berge et al., 2020). These approaches still used the PCA and KNN low-dimension system with statistical inference to draw an artificial trajectory. The trajectory feature of trajectory graphs created by BW all came from the nature of high-dimension data itself. In other words, if data points in the high-dimension space had no trajectory feature or agglomerated tightly with each other, no trajectory feature will be shown in the graph.

### Finding

The BW system revealed deep insight into human brain endothelial cell composition. Compared with the control sample hierarchical structure, the Alzheimer’s sample exhibited stem cell exhaustion and malfunctioning immune response. Surprisingly, the functional endothelial cell group in the Alzheimer’s sample still had an intact hierarchical structure while the overall lifespan of the Alzheimer’s sample almost doubled.

What is more, the BW system finding also supported that current sub-type markers of endothelial cells were not ideal, and positive senescent markers were highly possible only an illusion that was also reported from other recent research.

## Results

### Performance on raw matrix control samples

To assess the performance of BW and the PCA-based approach by Scanpy (Wolf et al., 2018), I conducted a comparison for each sample. The logged matrix was created by applying the log1p function to the raw matrix, while the raw matrix contained the original count matrix with all elements greater than or equal to 0. The PCA-based method employed only the logged matrix, which was included in their standard procedure by tutorial. For the BW-based clustering, the raw matrix was used. The projection of the PCA-based method on the BW method-based cluster visualization was obtained by using PCA-generated clusters to display on the BW method cluster graph.

In terms of visualization, the BW method-based cluster graph outperformed the PCA-based method in displaying cluster information (fig1, sup(fig1)) as clusters generated by PCA were better separated on the BW generated cluster graph. This was expected since BW visualization was based on the pre-generated BW clusters. On the other hand, the PCA-based method had to undergo a 2D partition before generating clusters. The cluster visualization was created before the clusters were formed in the PCA-based method. In comparison to the PCA-based method, BW clustering produced more detailed and realistic clustering, as the fading color trend was consistent throughout the graph (fig1 A3 B3 C3). It is not possible that elements for one cell type are just exactly located within a 2D visualization regular zone. Overall, the BW method showed high consistency with the PCA-generated general cell types for cell type clustering. Next, the BW clustering visualization method was used to create the trajectory graph for samples (fig2).

**Fig 1.**
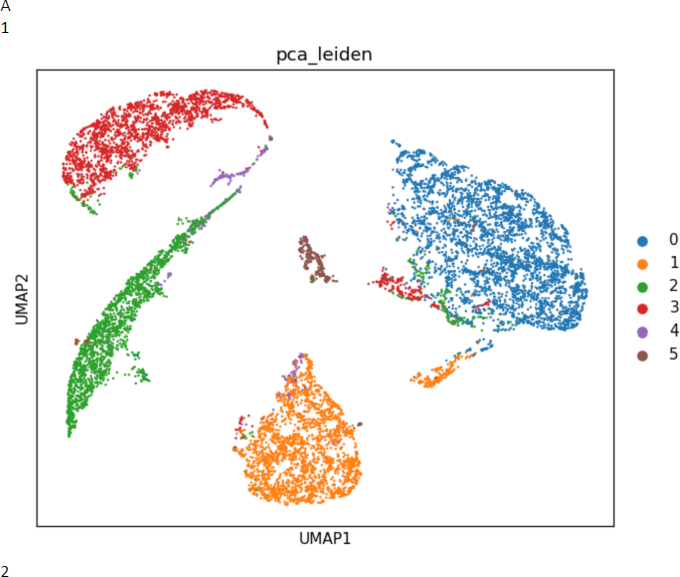
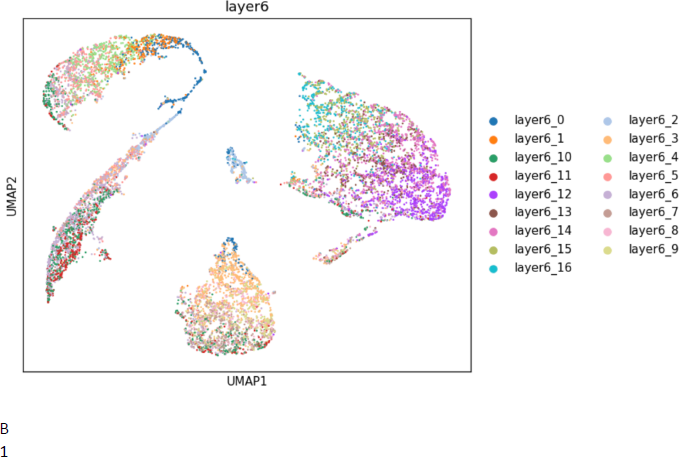
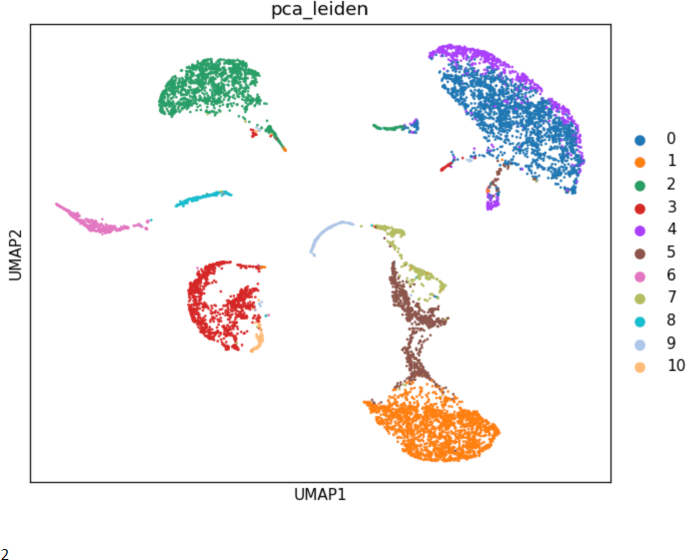
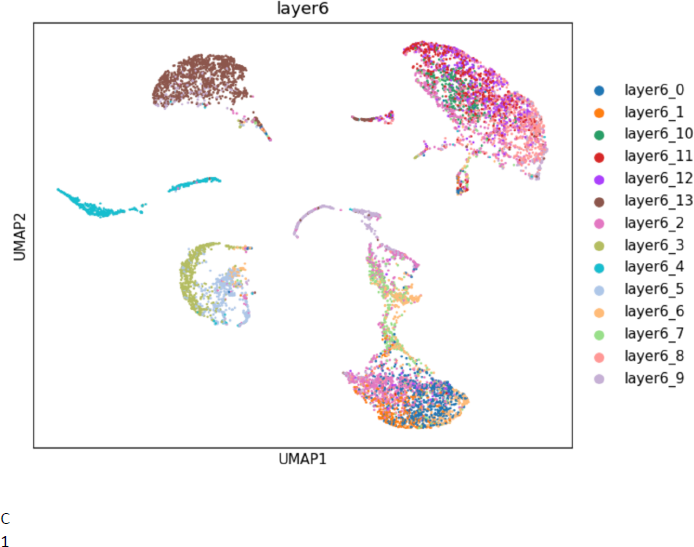
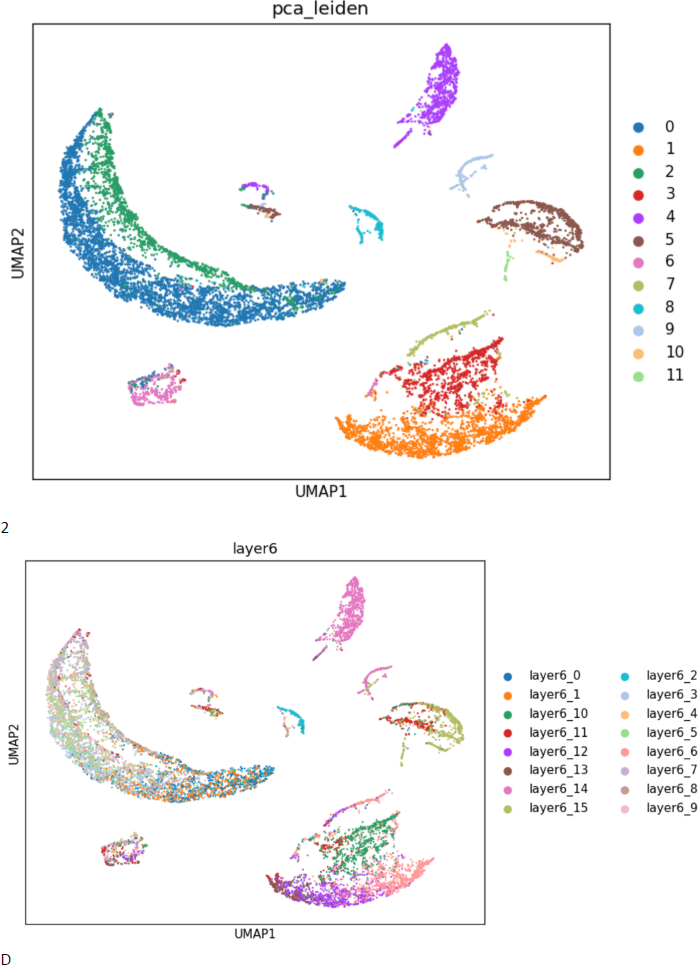
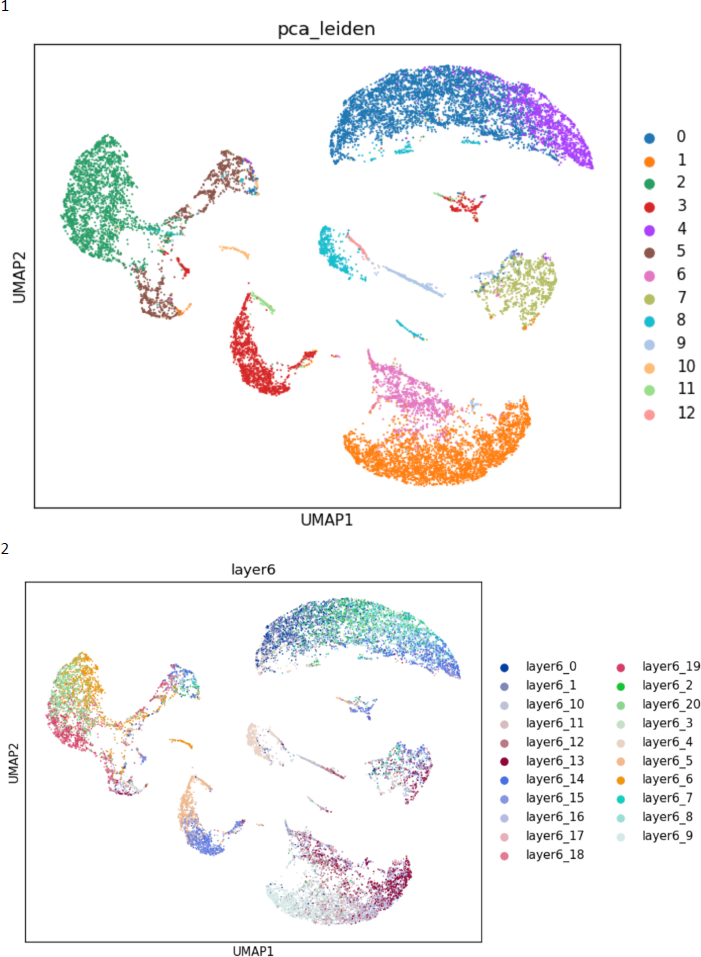
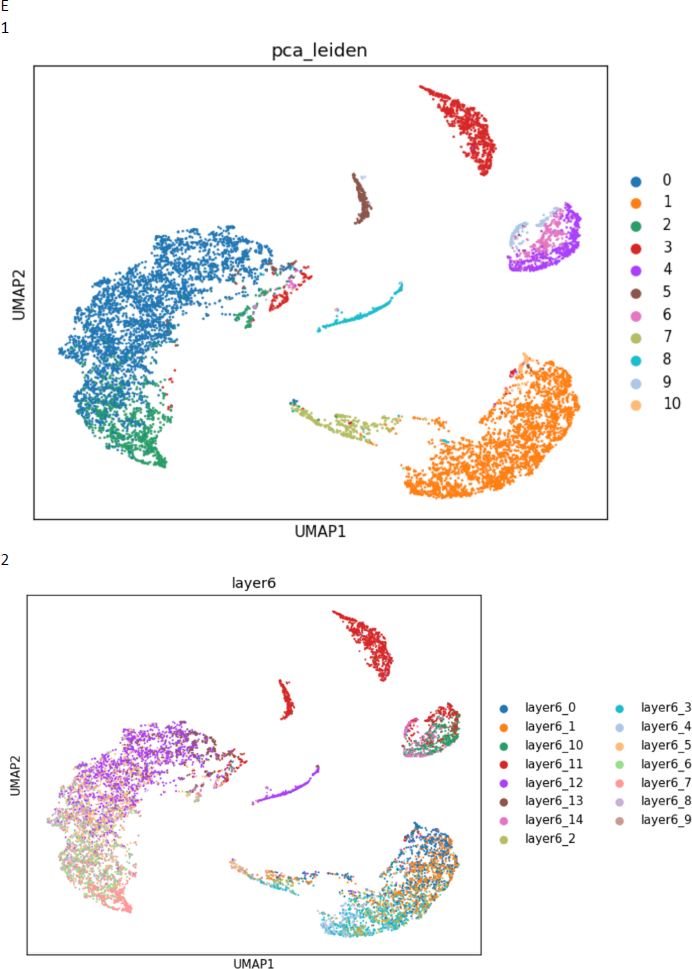
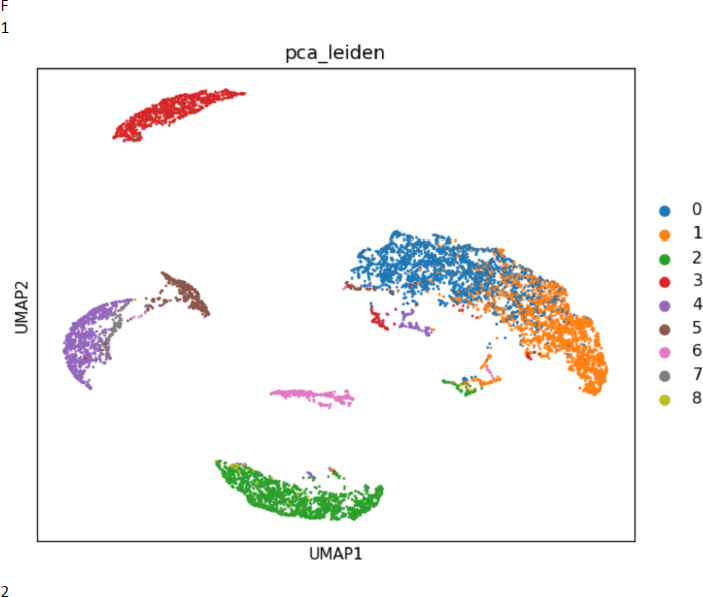
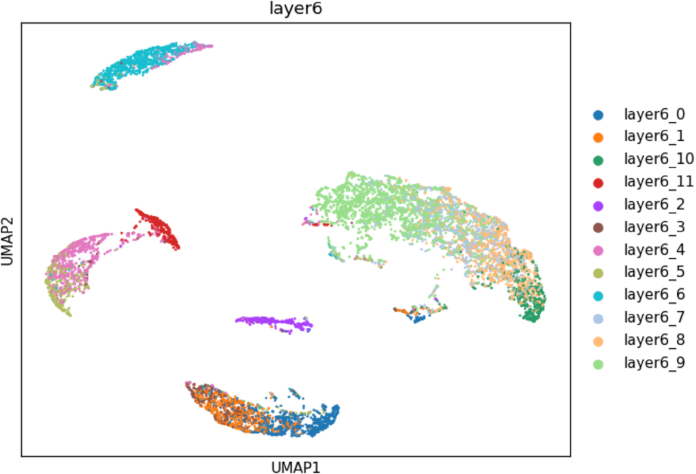
Legend: Cluster graph (1): PCA-based results projection on BW method-based cluster visualization. (2): BW method-based clusters and visualization of layer 6. (A, B, C, D, E, F): sample control1, control2, control3, AD1, AD2, AD3.

**Fig 2.**
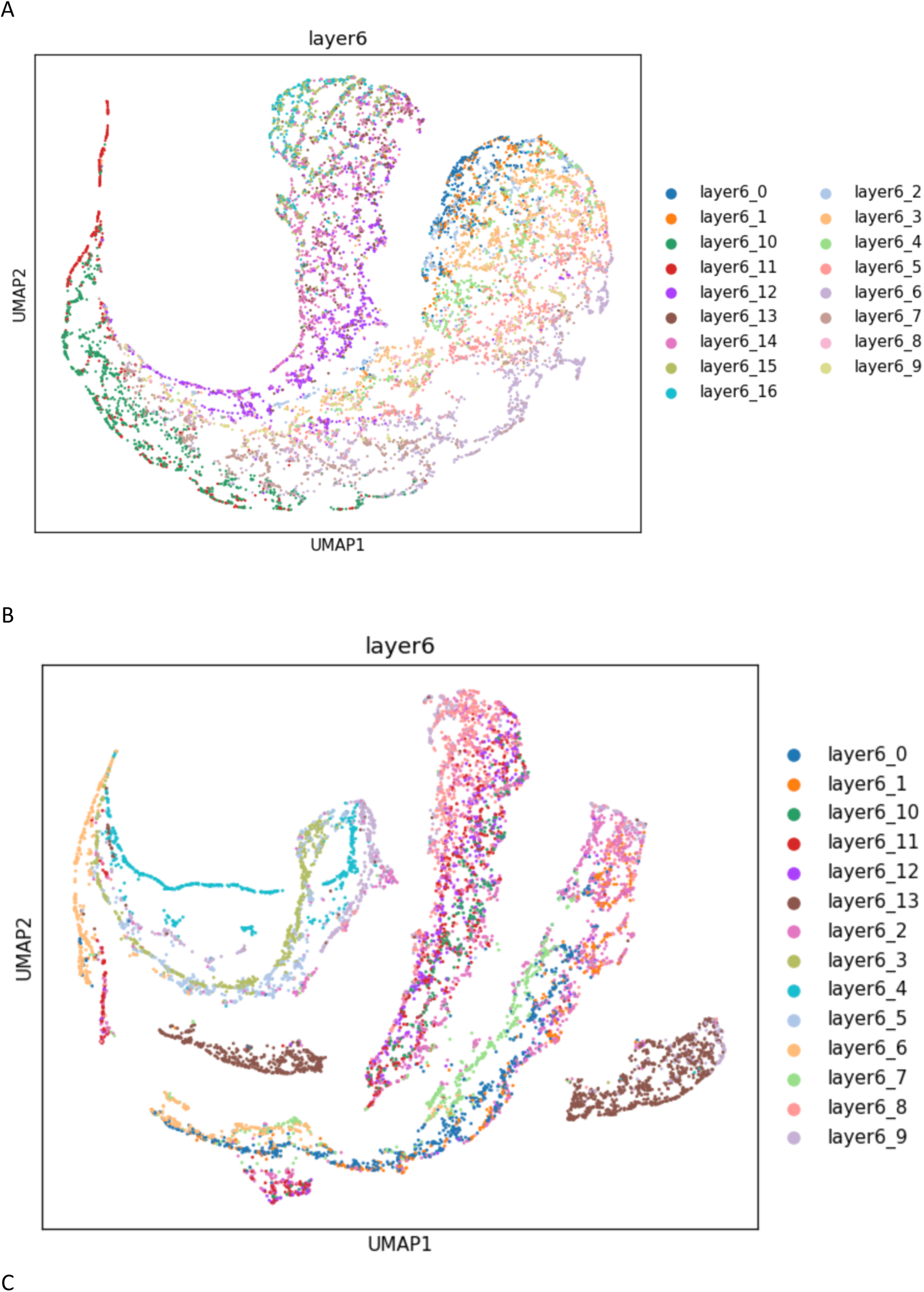
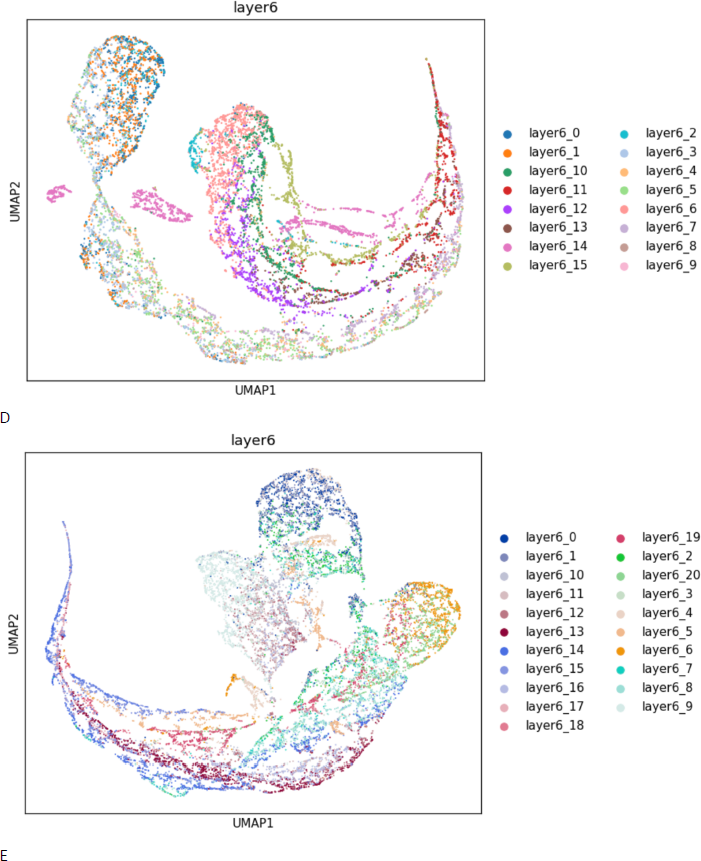
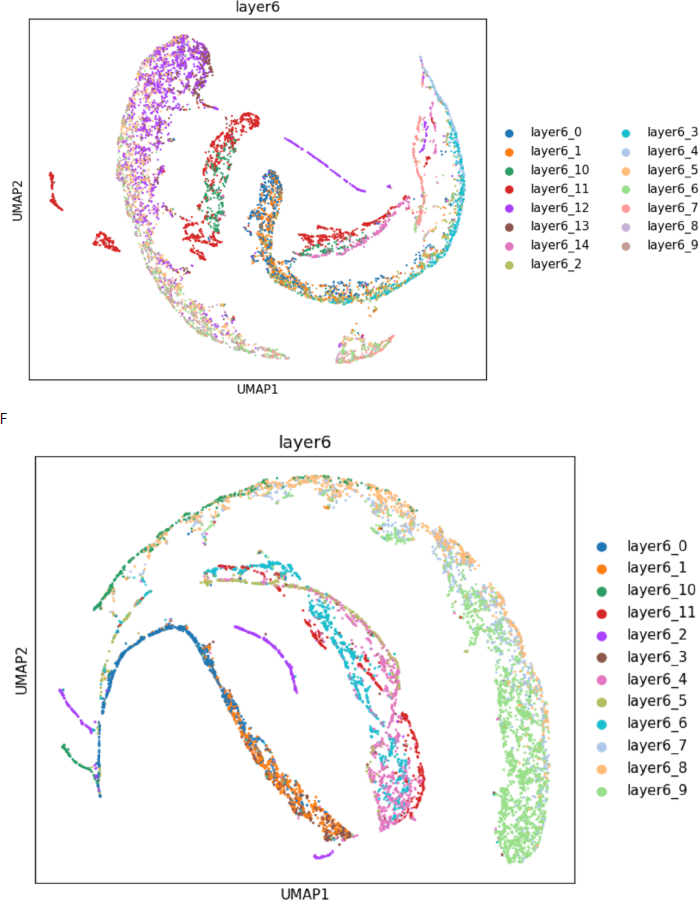
Legend: global trajectory graphs. (A, B, C): Sample control1, control2, control3, AD1, AD2, AD3.

The global trajectory would be very useful for researchers who just want brief and distinguishable graphs to show the location relationship between large general cell types (fig2). The results from different projection layers seemed not to have a significant difference. Theoretically, the higher the projection layer was, the accuracy of the graph would slightly decrease as the lower layer level has more category cores. For the redundancy of caution, I would recommend using the lowest layer while the clusters in that layer do not exceed 500 because the more category cores are, the high instability of UMAP reduction would be. The balance between accuracy and instability is always the art of data science.

In summary, the results on raw matrix data supported that the BW clustering and visualization method outperformed the PCA-based system. The BW visualization only tries to show the true shape of high-dimensional data without any 2D partition. Leveling up the projection layer of the BW visualization graph did not show a significant difference.

### Trajectory practical explanation and guidance

#### Stem cell, aging, and differentiation feature

The head part of a cluster’s trajectory should be the earliest stage of cell age. The stem cell branch should have a dominant proportion within the head part of each cluster. Compared with functional cells, the gene expression lifespan of the stem cell branch should be shorter (Fig3 A). When aging stage changes became dominant, the cluster will show a clear color segmentation pattern which would be very easy to identify (Fig3 B).

**Fig 3.**
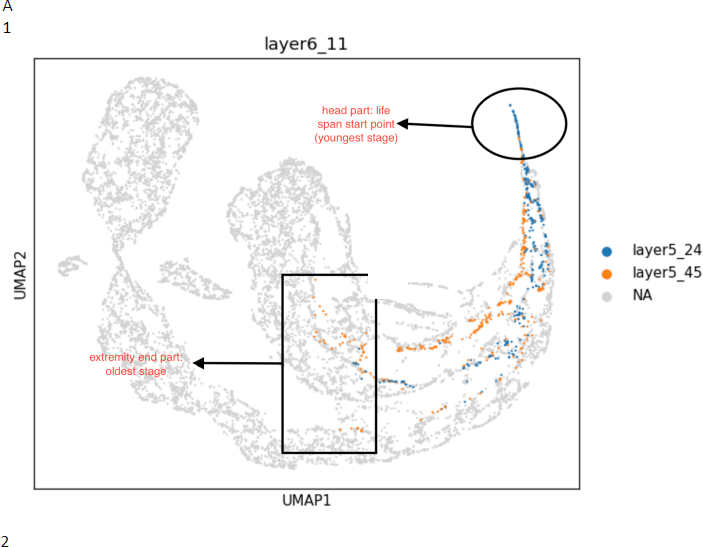
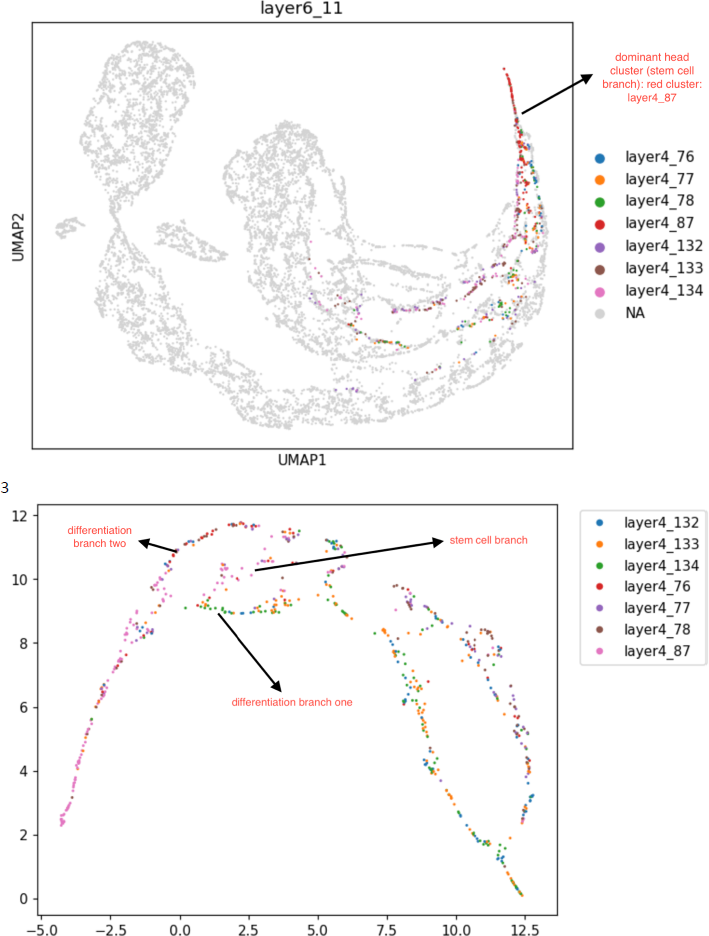
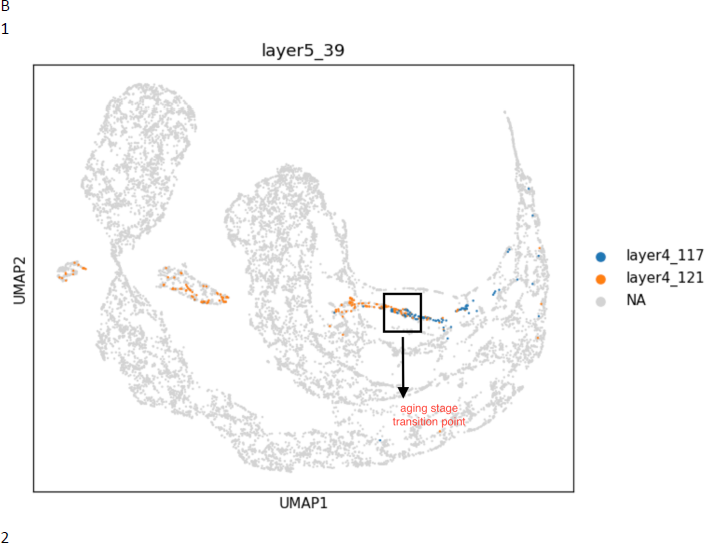
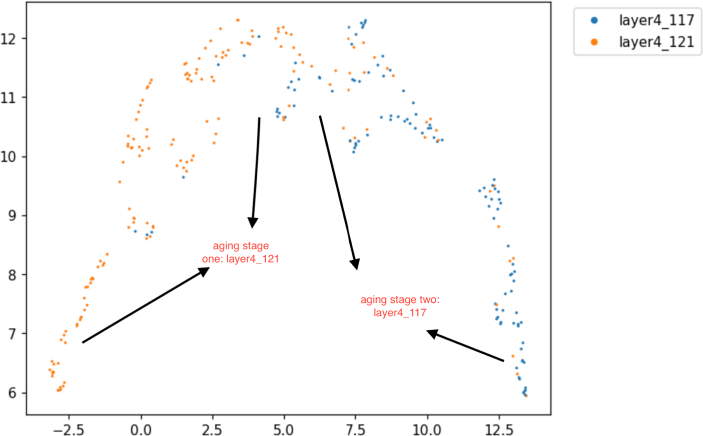
Legend: trajectory graphs (A): (1) global trajectory graph of control3 sample layer6_11 cluster descending to the layer5 level. (2): descending to the layer4 level of (1). (3): partial trajectory graph of control3 sample layer6_11 cluster descending to the layer4 level.

#### Escaping effects (gene expression instability)

**Schematic diagram 1:**
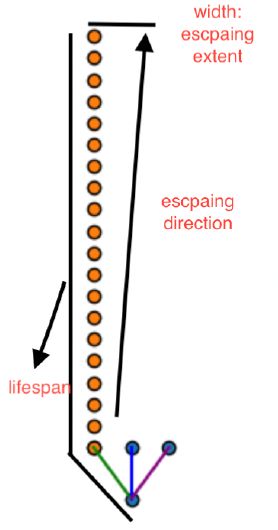
The arrow indicates the escaping direction.

When cells escape from a differentiation branch, they do not move perpendicularly. Similar to an object thrown from a moving train or airplane, the escaped cells will not drop vertically to the ground. Instead, they will move close to the orbit for a long distance. The strength of the escaping effect, as well as the distance between the escaping nodes on the cell state line, will determine how clear the cell state change is. When escaping effects became dominant, this would normally result in a totally overlapping color pattern (Fig4 A). In this case, it is more likely that the cluster only has one single differentiation branch which means it does not have sub-types.

#### Examples of different types of sub-clusters

Escaping effects:(Fig4 A)

Aging stage changes: (Fig4 B)

Cell development changes, subtypes, and differentiation: (Fig4 C D)

**Fig 4.**
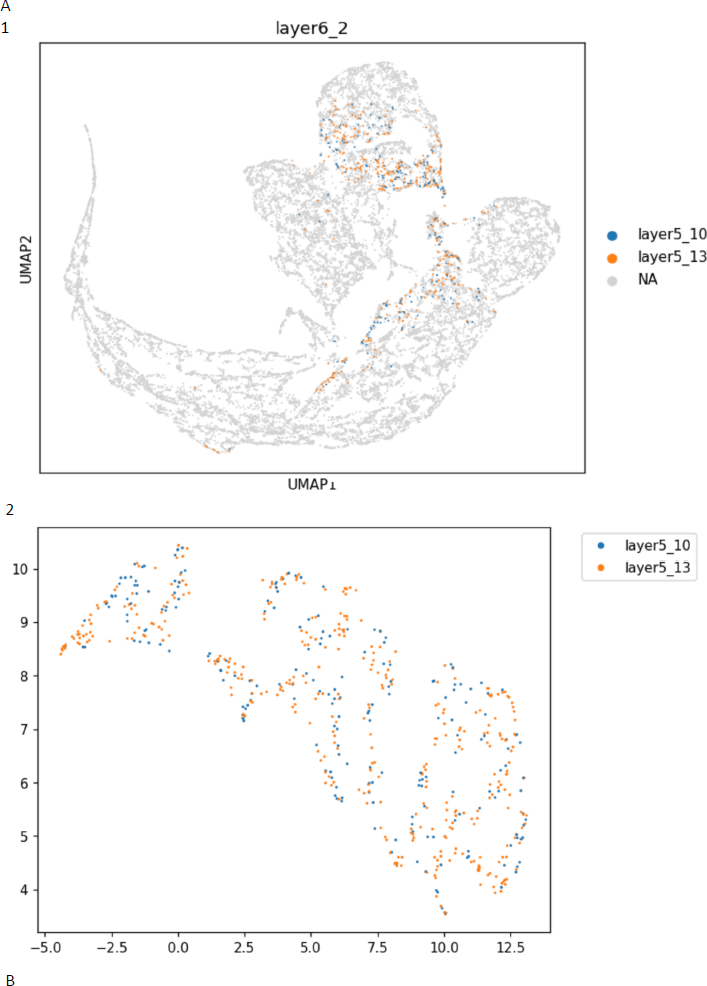
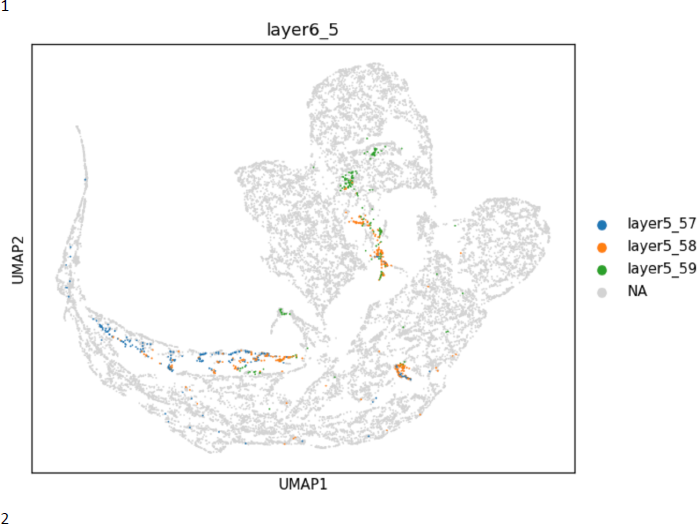
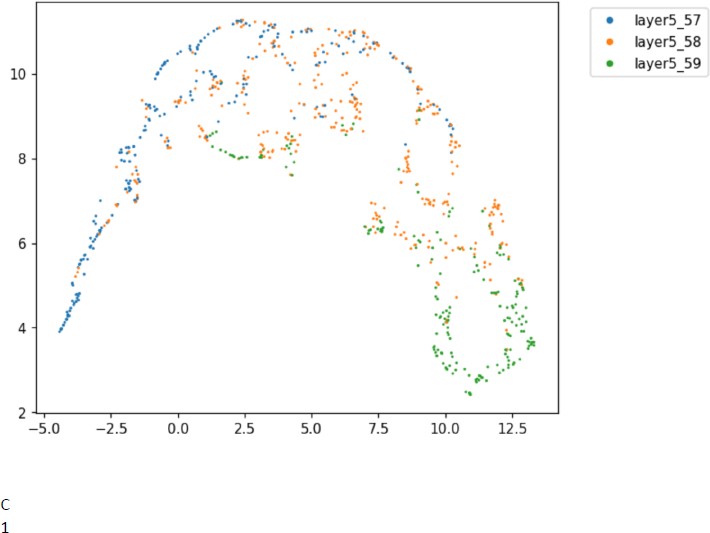
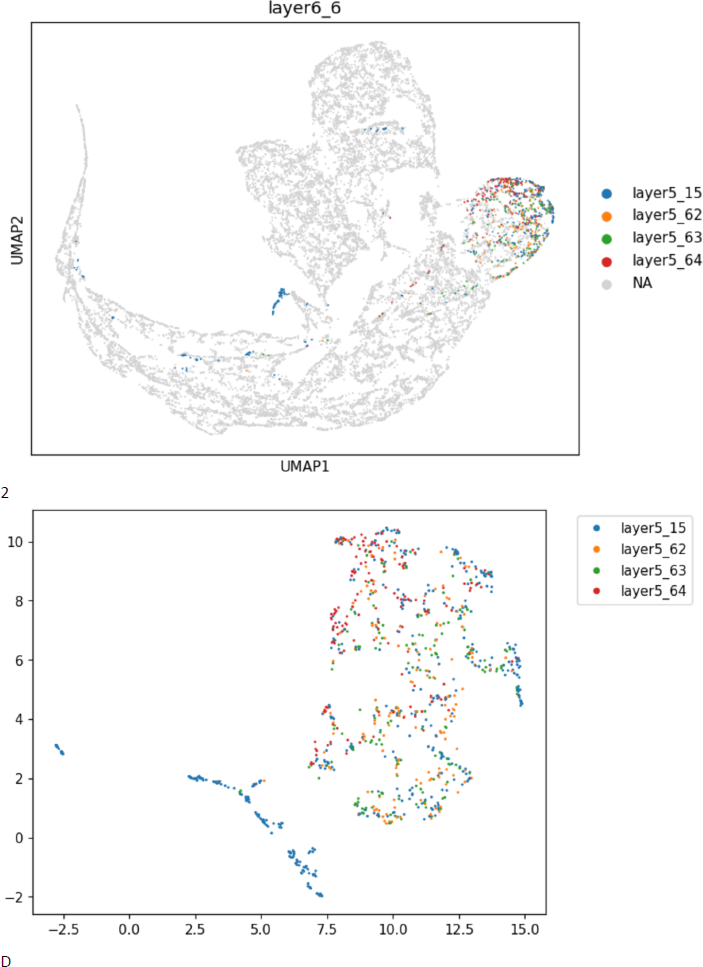
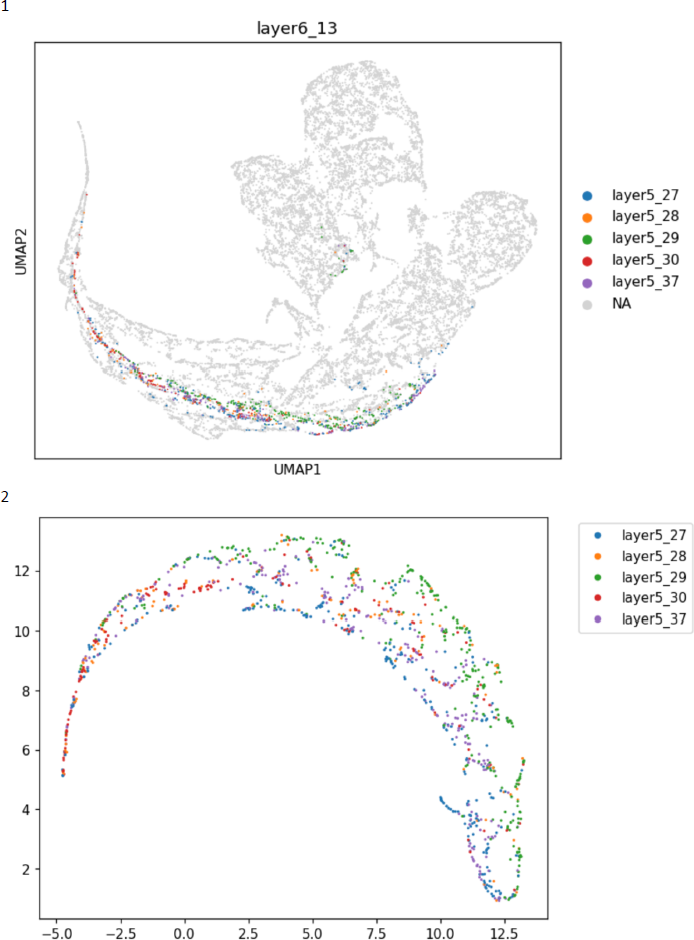
Legend: Trajectory graphs (1, 2): global, partial trajectory. (A, B, C, D): cluster layer6_2, cluster layer6_5, cluster layer6_6, cluster layer6_13 from AD1 sample.

### Endothelial cell analysis

After annotation by endothelial marker (’FLT1’, ’CLDN5’), 5821 endothelial cells were found from the control3 sample (total cell number: 12142) achieving a 0.4794103113160929 ratio of endothelial cells. AD1 sample showed 6716 endothelial cells with a total cell number of 18839 generating a relatively lower ratio: 0.3564945060778173. Other samples’ ratios were too far behind AD1 and control3. Thus, the following analysis only used the AD1 sample and control3 sample.

First, the mainstream PCA-based secondary PCA partition was used to test the widely accepted endothelial cell subtype marker: {capillary: ’SLC16A1’, arterial: ’BMX’, venous: ’NR2F2’, arterial: ’VEGFC’}. With all gene dimensions, the secondary PCA partition result (Fig5 A) generally showed a big mass which means no significant sub-clusters. The common markers distribution also did not have any regular pattern within this approach as well (Fig5 A2). To thoroughly analyze PCA based method’s approachability, I only used four marker genes as dimensions to run the PCA-based secondary partition again. The result became even worse (Fig5 B). Due to the advantages of hierarchical structure, the BW system does not need to run further analysis to reach sub-clusters. The result showed that although BW sub-clusters can be divided into three major clusters, there was no generally distinct pattern of four sub-type markers as well. The finding matched recent other studies on those markers (Chen et al., 2020; Ximerakis et al., 2023).

**Fig 5.**
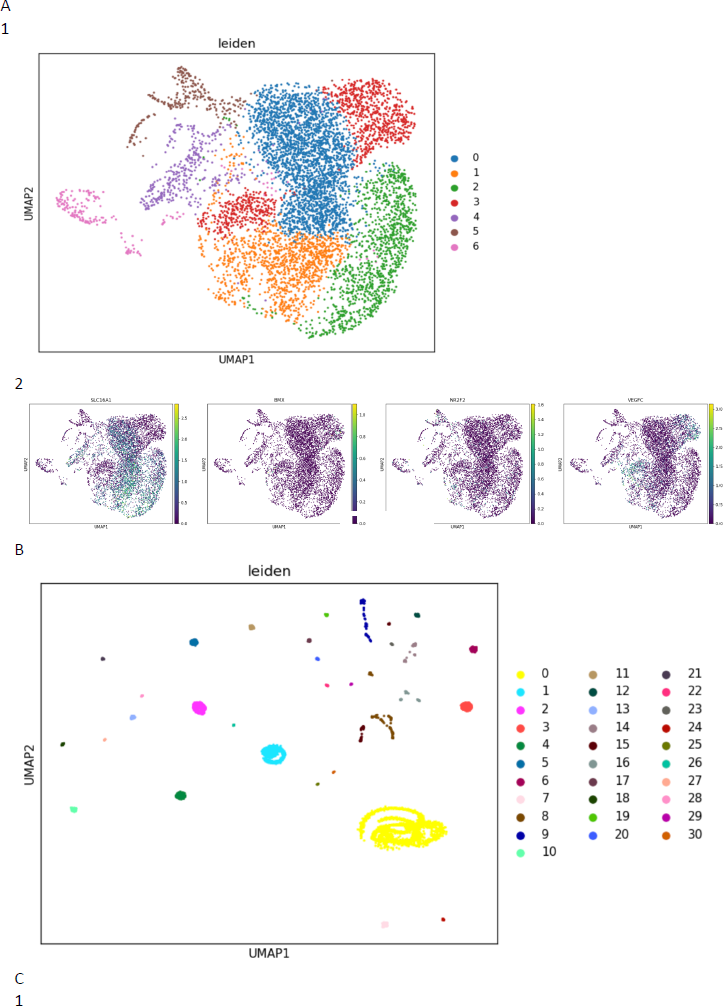
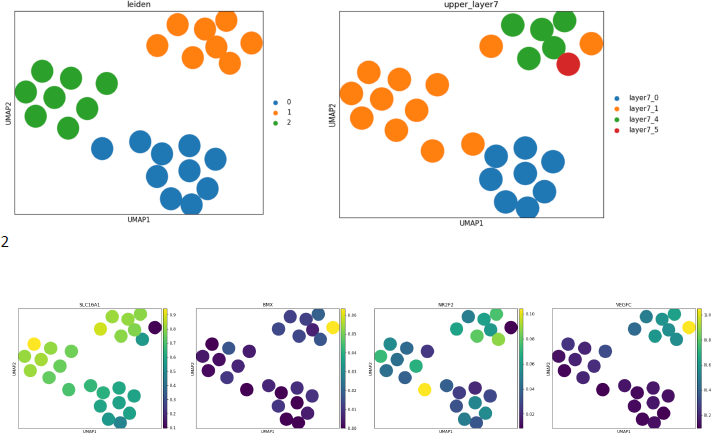
>Legend: PCA umap reduction graphs (A): PCA secondary analysis on endothelial cells from sample AD1 with all gene dimensions. (B) PCA secondary analysis on endothelial cells from sample AD1 with only four sub-type marker dimensions. (C): PCA secondary analysis on the BW layer5 endothelial sub-clusters from sample AD1 with only four sub-type marker dimensions.

The BW trajectory graphs showed a high consistency of endothelial sub-types between the two samples (Fig6 A) on the layer7 level. AD1 sample exhibited a unique final aging stage: layer7_0 (Fig6 A1). This unique aging stage had a lower level expression of all four sub-type markers (Fig5 C2). Combing with the marker expression and location of the unique aging stage on the trajectory graph, it was not hard to infer that this would be an Alzheimer’s disease special senescent endothelial cell group. Both two samples shared a common next-level sub-type structure, they both had a functional cell sub-type (AD1:layer7_1, control3: layer7_0), one head part dominant cluster (AD1:layer7_5, control3: layer7_3), secondly head part dominant cluster (AD1: layer7_4, control3: layer7_2). To further analyze the difference between the control and disease samples, I compared all three sub-type lower-level structures (Fig6 B C D E). The functional cell sub-type between the two samples shared an extremely similar inner structure on the layer6 level (Fig6 B), they both had a distinct branch sub-type in red color (AD1:layer6_8, control3: layer6_3) and other three overlapping sub-clusters which were showing escaping effects and suspected to be three commonly known sub-types: capillary, venous, and arterial. Even after diving into layer5 level, those two red clusters still shared a very similar high-dimensional shape (Fig6 C). The second head part dominant clusters also had a similar shape but the AD1 sample did not have a layer6 sub-type indicating one branch that existed in the control sample vanished in the disease group (Fig6 D). The cluster that had the biggest stem cell feature in the AD1 sample changed dramatically compared with its counterpart in the control sample (Fig6 E).

**Fig 6.**
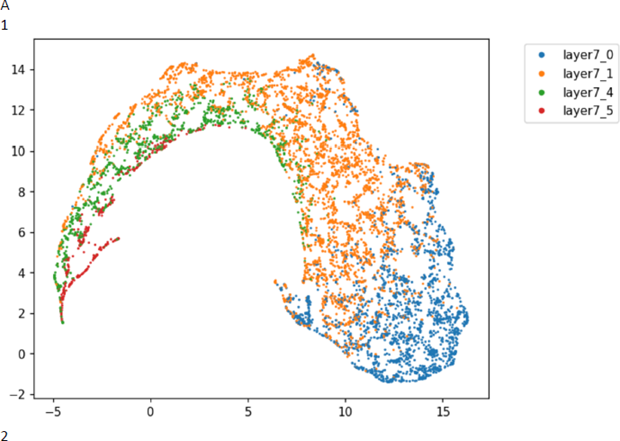
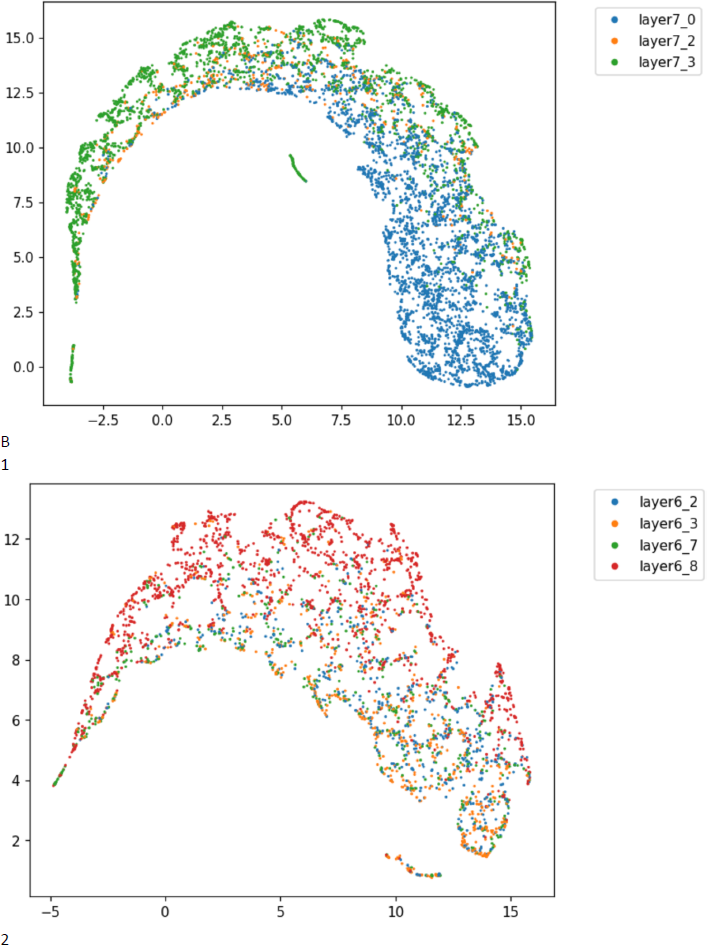
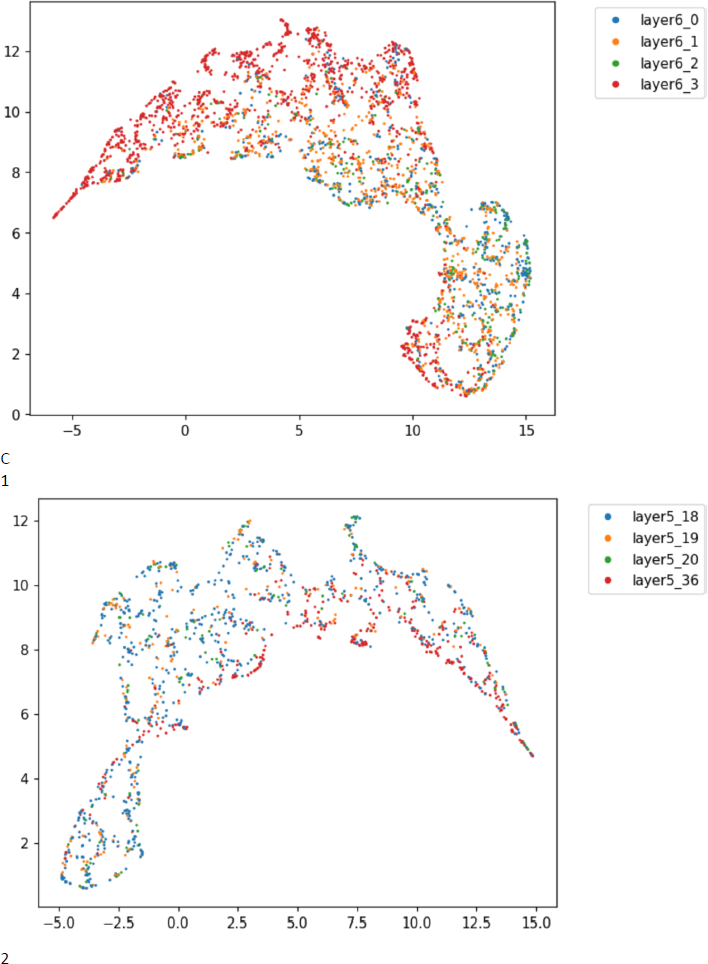
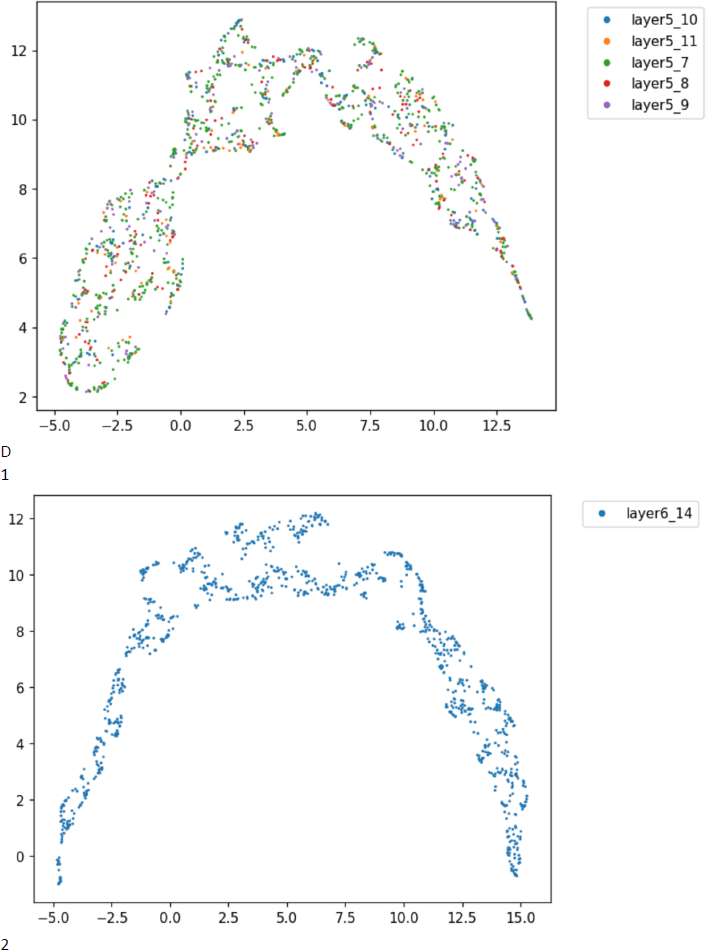
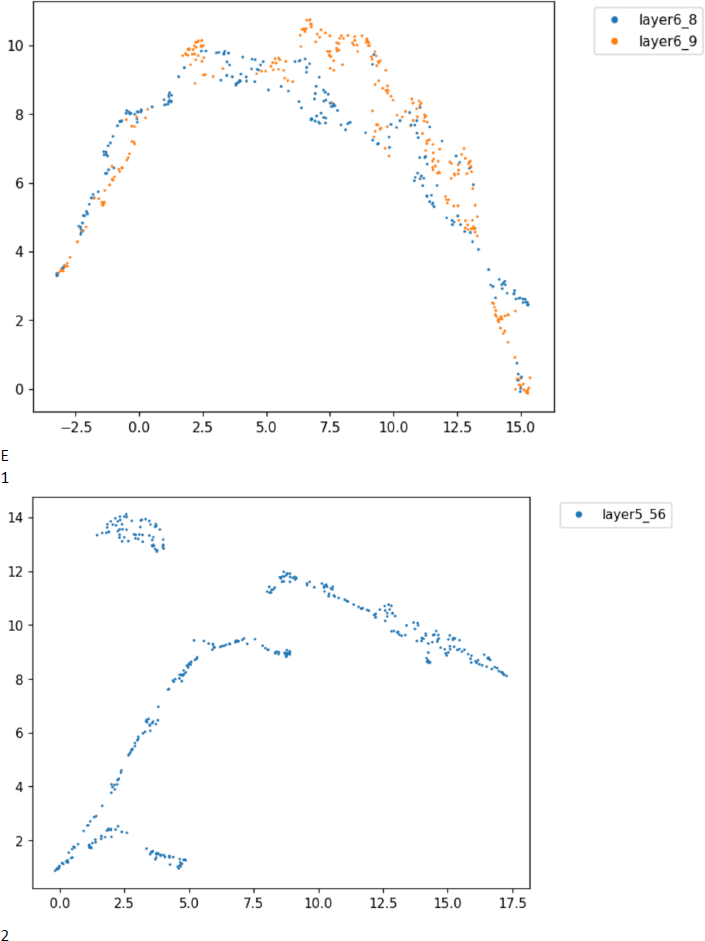
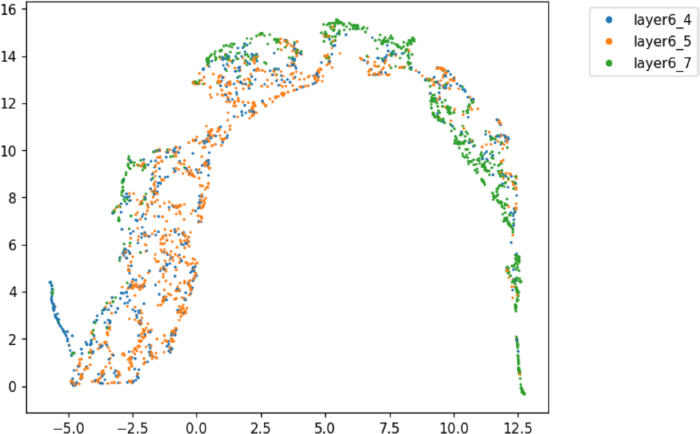
Legend: partial trajectory graphs (A, B, C, D, E): endothelial cell, functional cell sub-type, red distinct clusters in B, second head part dominant sub-type, head part dominant sub-type. (1,2): AD1 sample, control3.

In summary, The BW system unveiled a comprehensive, accurate, deep, and consistent hierarchical structure of endothelial cells which made the comparison of hierarchical structure between control and disease groups available. It showed that human brain endothelial cells had three sub-types (two sub-type with stem cell features, and one functional cell type) on the secondary type level. On the third type level (layer6) and fourth type level (layer5), hierarchical structures made by the BW system unveiled the functional endothelial has four sub-types where one was distinct, and the other three were highly similar. Linking all levels’ findings, the Alzheimer’s disease sample had a special senescence cell group and two distorted stem cell feature groups while the functional cell group was exactly the same as the control sample suggesting that stem cell exhaustion and a compromised immune system could be potential factors that contribute to Alzheimer’s disease. What is more, the three highly similar sub-types for functional endothelial cells appeared to be capillary, venous, and arterial as their markers generated by Scanpy rank_gene_group were highly related to shear stress indicating that capillary, venous, and arterial are not real differentiation procedure but an adaption to the shear stress.

### Secondary analysis

In order to better understand the cell aging state changes, cell gene expression lifespan, and gene expression instability. A secondary analysis is required. To perform a secondary analysis, the target could be either the same thing between different samples or singular branches. For example, a comparison between T cells in different samples is allowed. A comparison between T cells in one sample and CD34+ T cells in another sample is not allowed. The sub-type or sub-cluster which only had one differentiation branch (singular branch) would be always available to compare.

Both endothelial cells in the AD1 sample and control3 sample had 8 eight aging stages (Fig7 A) after the secondary analysis partition. To validate how successful the secondary partition and BW system were, the top 40 genes generated by Scanpy rank_gene_group of the first aging stage cluster (Fig7 B) were inspected. The consistency of the general decreasing pattern was found in both samples. Thus, the secondary analysis and the BW system’s biological meaning were both validated.

**Fig 7.**
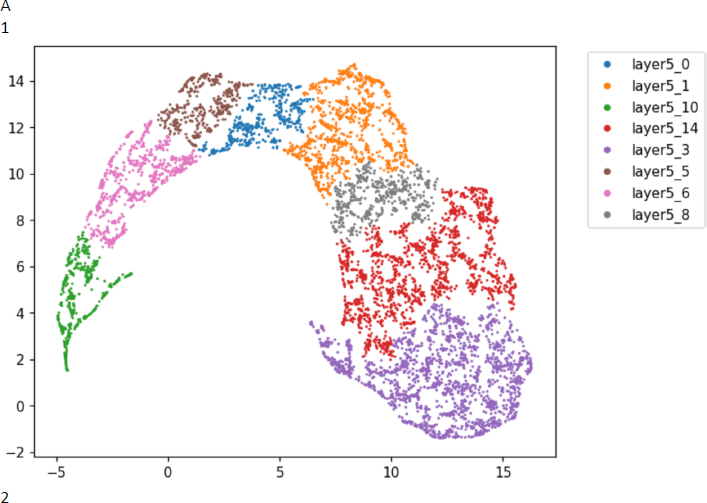
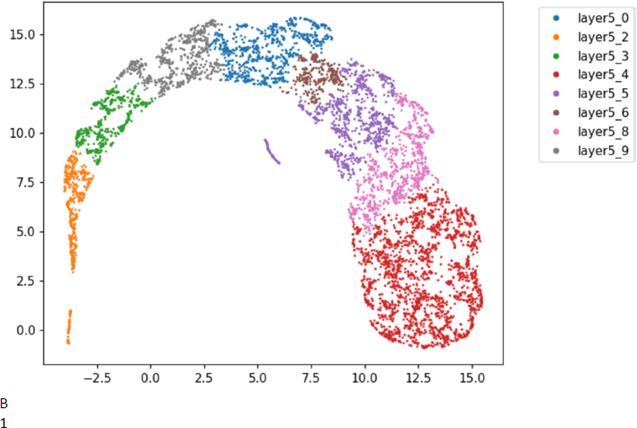
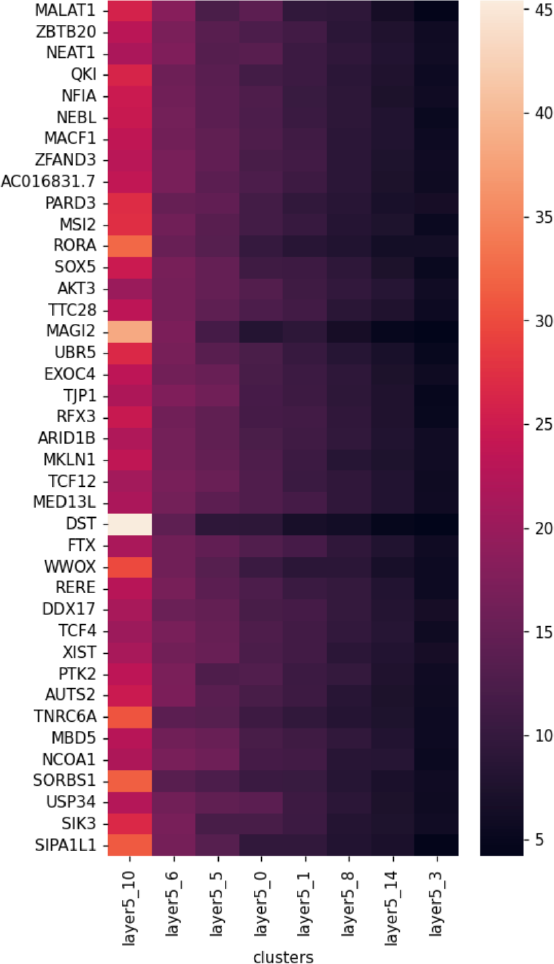
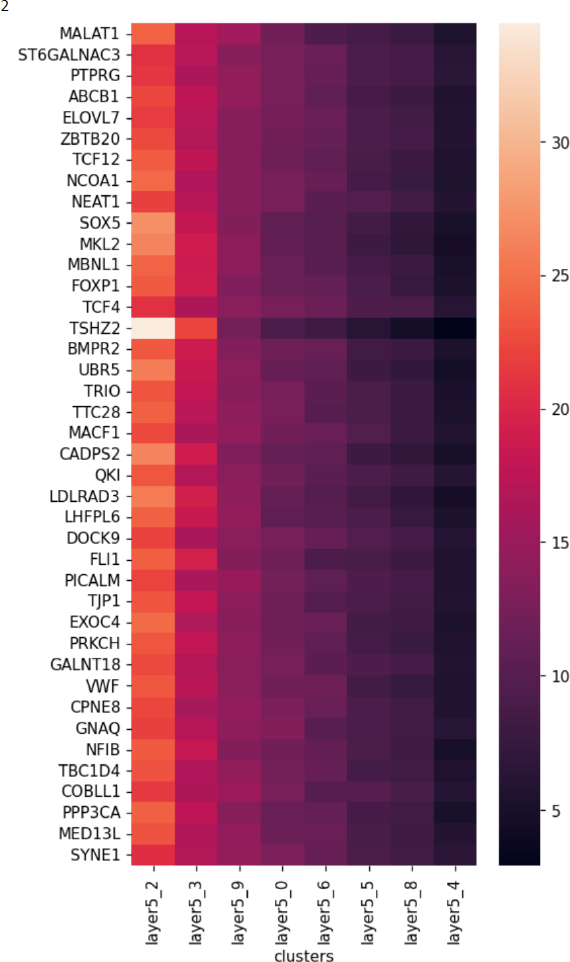
Legend: (A) endothelial cell partial trajectory graphs with secondary analysis cluster partition. (B) heatmaps of the top 40 marker genes of the first aging stage secondary analysis cluster made from Scanpy rank_gene_group function. (1,2): AD1, control3.

The secondary analysis aims to get three components: gene expression instability extent, gene expression lifespan, and each aging stage’s markers. From gene expression instability graphs (Fig8 A), the Alzheimer’s sample showed a relatively low gene expression instability in the early stages whereas a higher instability pattern in the middle and late aging stages. The lifespan of the Alzheimer’s sample was nearly twice of the control3 sample. The secondary analysis results matched the primary hierarchical structure comparison as the less number of stem cells would apparently result in a lower instability in the early stage and abnormal senescent cells which immune cells failed to remove can definitely expand lifespan and expression instability in the late stages.

**Fig 8.**
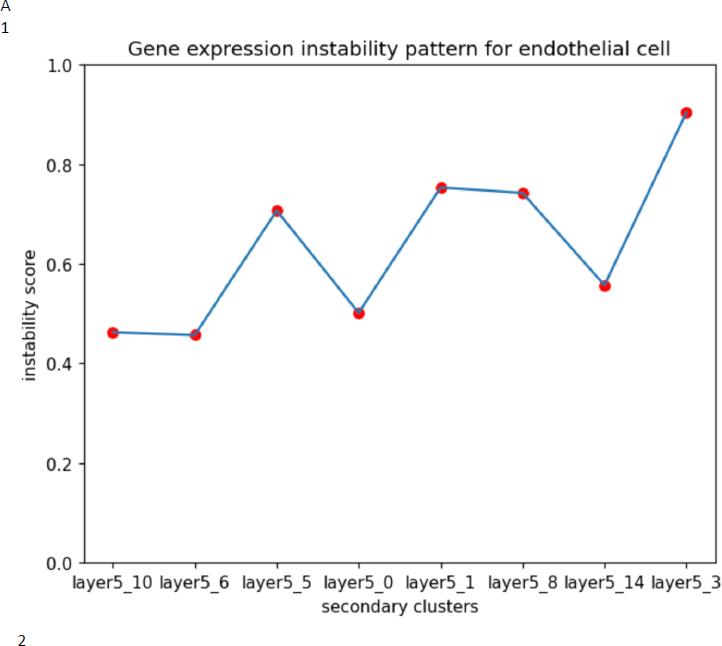
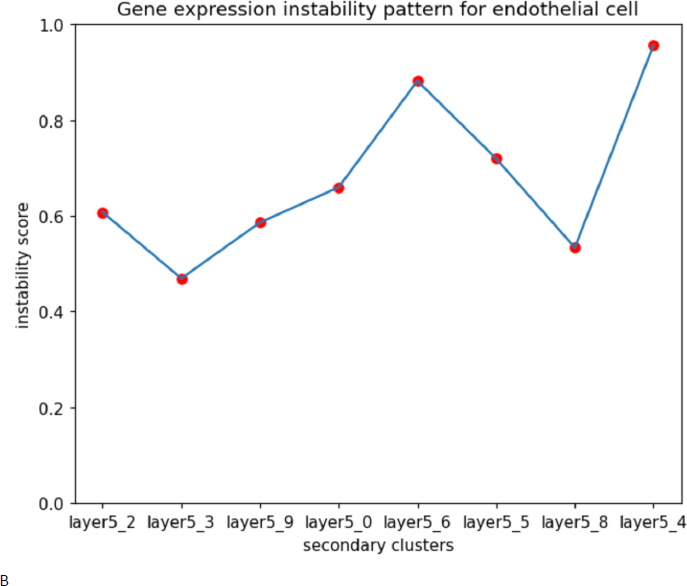
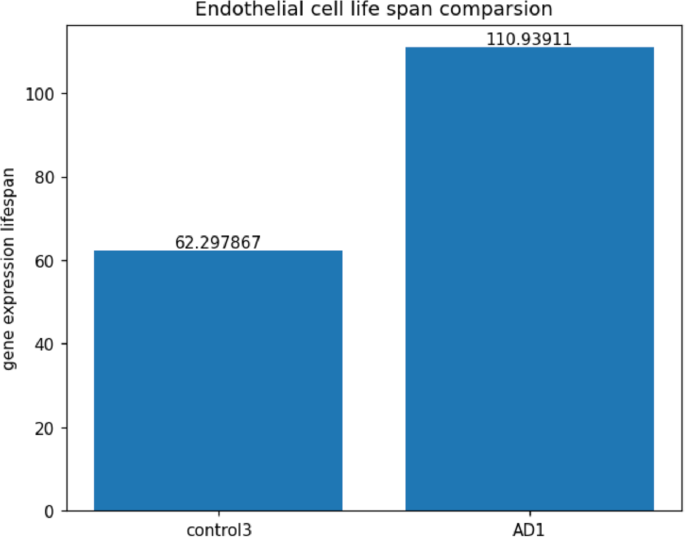
Legend: (A): gene expression instability graph prolonging the aging process (1,2): AD1, control3. (B): lifespan comparison between AD1 and control3.

The secondary analysis also gave the unique aging stage markers for each aging stage. The BW secondary analysis marker finding algorithm was different from the general type marker finding algorithm as it applied a multiple-group comparison strategy while general marker finding algorithms like Scanpy rank_gene_group usually adopted a two-group comparison. An intriguing fact was that along the cell differentiation branch, as cells become older, the number of positive markers hugely shrank (sup(fig2)). The specificity of positive markers also decreased dramatically. In other words, most genes became more and more silent with the progression of aging or senescence. Positive markers between each cell state stage showed high heterogeneity. Perhaps, this is probably the reason why aging or senescence markers are so hard to find and replicable because markers for each stage are quite different. Combined with the fact that there was no available hierarchy clustering method and corresponding marker-finding algorithm on high-dimensional large data, it is possible that the so-called aging marker which can distinguish the senescent cell and the non-senescent cell does not actually exist as aging or senescence is a linear or continuous process. In fact, the debate of senescence markers widely existed as the test on those markers was not positive at all (Ximerakis et al., 2023). This trajectory graph-based secondary analysis could be a better choice for aging studies.

### Some tests on the purged matrix

To investigate how the number of dimensions impacts the performance of BW and potential solutions for better results integration ability of multiple samples, a purged function was utilized to normalize and filter the dimensions. Specifically, a dimension was deemed valid if the difference between its maximum and minimum values was between 3 and 350 across all cells. Any non-zero values were converted into percentages and scaled by the dimension’s average. In this study, the PCA-based method utilized the same dimensions and logged matrix, while all BW-based graphs were generated using layer3 as the projection layer, and the BW cluster graph was based on the logged purged matrix.

The purge function in BW can effectively reduce the magnitude of escaping effects and shift the observation site from outside to inside, resulting in reduced noise but increased data dispersion which leads to unclear differentiation start points. This observation was validated through practical experimentation (see figS3). While the BW cluster graphs clearly showed that many of the same clusters were separated into parallel shapes (figS3, B, E, F), the logged global trajectory graphs became increasingly unstable as the data shape became more dispersed. As a result, the clustering patterns in figS3 (B, E, F) started to deviate slightly from the PCA-based clustering approach, whereas figS3 (A, C, D) remained tightly aligned with PCA-based clustering. It is difficult to determine which approach is better in this case, as they may be biased in different directions. However, comparing only the purged global trajectory graphs shows that BW outperformed PCA-based clustering slightly when the data was purged (figS3 B5, E5, F5 compared to fig9 A1, A2, A3). The difference between figS3 (A, C, D) and figS3 (B, E, F) can be attributed to the preserved number of dimensions. Specifically, the purged matrices of control1, control3, and AD1 contained 5336, 7664, and 6282 genes, while the remaining samples had only around 3000 genes in their purged matrices.

**Fig 9.**
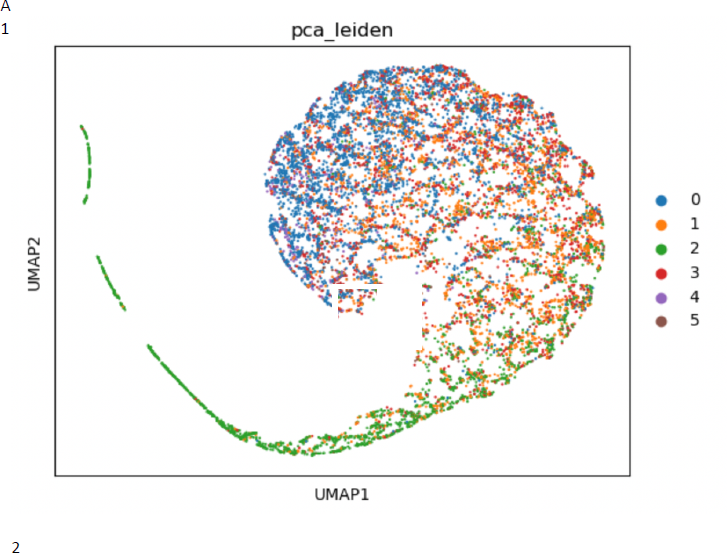
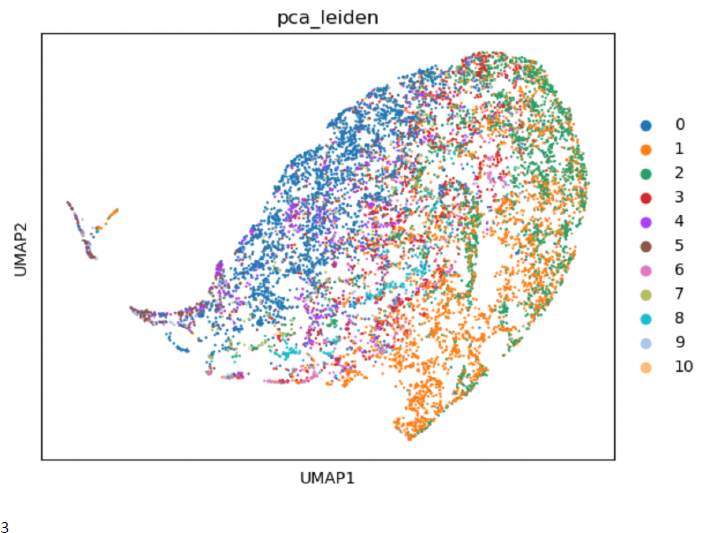
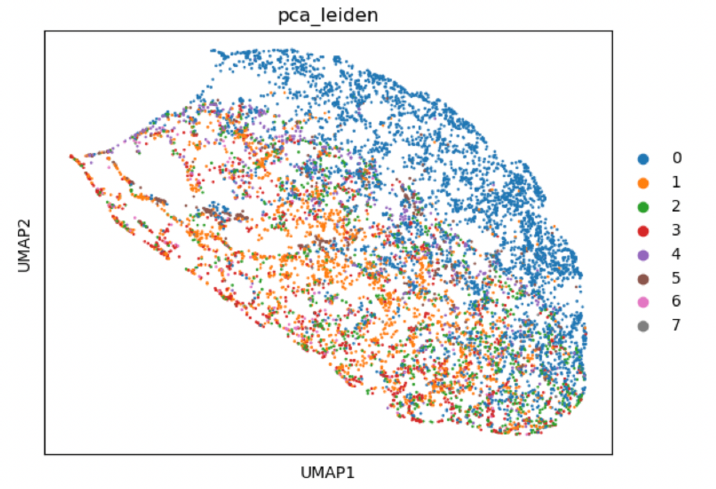
Legend: PCA-based clusters’ projections on corresponding BW global trajectory graphs. (A, B, C): Control2, AD2, and AD3 samples respectively.

### Speed time and resources

After conducting experiments, it was found that while the speed of the BW clustering method may not be super-fast, it is still a practical speed given the limited resources available. What is more, with the Apple AMX-optimized Numpy module, all the running times in Table 6 plummeted to one-tenth (**the speed surged 10 times**).

**Table 6:** Running time table for BW clustering.

| Sample | Running time | Number of cells | Number of genes | Resource |
| --- | --- | --- | --- | --- |
| control 1 | 39 mins and 12 seconds | 12142 | 33538 | m1 chip MacBook air with 16GB memory |
| control 2 | 32 mins and 41 seconds | 11048 | 33538 | m1 chip MacBook air with 16GB memory |
| control 3 | 27 mins 12 seconds | 10039 | 33538 | m1 chip MacBook air with 16GB memory |
| AD1 | 110 mins 20 seconds | 18839 | 33538 | m1 chip MacBook air with 16GB memory |
| AD2 | 27 mins 57 seconds | 10600 | 33538 | m1 chip MacBook air with 16GB memory |
| AD3 | 33 mins 24 seconds | 9101 | 33538 | m1 chip MacBook air with 16GB memory |
| control 1 | 8mins and 45 seconds | 12142 | 5336 | m1 chip MacBook air with 16GB memory |
| control 2 | 5mins and 9 seconds | 11048 | 2753 | m1 chip MacBook air with 16GB memory |
| control 3 | 7mins and 53 seconds | 10039 | 7664 | m1 chip MacBook air with 16GB memory |
| AD1 | 24mins and 7 seconds | 18839 | 6282 | m1 chip MacBook air with 16GB memory |
| AD2 | 5mins and 35 seconds | 10600 | 3846 | m1 chip MacBook air with 16GB memory |
| AD3 | 3mins and 59 seconds | 9101 | 3151 | m1 chip MacBook air with 16GB memory |

However, there is still room for improvement in terms of speed optimization. One potential solution is to consult with a computer scientist to optimize the functions and compile them to enhance speed. Moreover, there is ample scope to refine the current parameters set, which could lead to a significant reduction in running time. By implementing these improvements, the BW clustering method can become even more efficient and accessible to a wider range of researchers.

## Conclusion

The BW clustering in high-dimensional data analysis allows for unprecedented insights into single-cell RNA sequence data. Firstly, it made a practically available hierarchical structure of high-dimensional large data. Secondly, the visualization system built on that structure can give researchers helpful guidance on what biological meaning and biological relationships are for clusters within layers. Finally, the secondary analysis system could present enlightenment for research on the cell state or development. While it currently only was used on very limited resources, further development and optimization are necessary to fully realize its potential. Machine learning and statistical approaches can be incorporated into the system to adapt to different sample sizes and qualities, greatly improving both the running time and precision of the layer levels. With continued refinement, BW has the potential to revolutionize how biological and biomedicine scientists approach biological high-dimensional large data analysis, ultimately leading to a deeper understanding of cellular biology.

## Data availability

All the matrix data were from NCBI Gene Expression Omnibus (GEO) under accession code GSE163577 (Yang et al., 2022).

Control samples id: GSM4982083, GSM4982085, GSM4982086 AD samples id: GSM4982084, GSM4982089, GSM4982090

## Disclosure

Please note that the content of this manuscript has been filed into a patent application.

