## Supplementary material for "A brand-new clustering method and analysis system designed for revealing the truth of the high-dimension large data deciphered the complex composition structure of human brain endothelial cells from single-cell RNA sequence data": sup figures

### Supplementary figures

Fig1

A

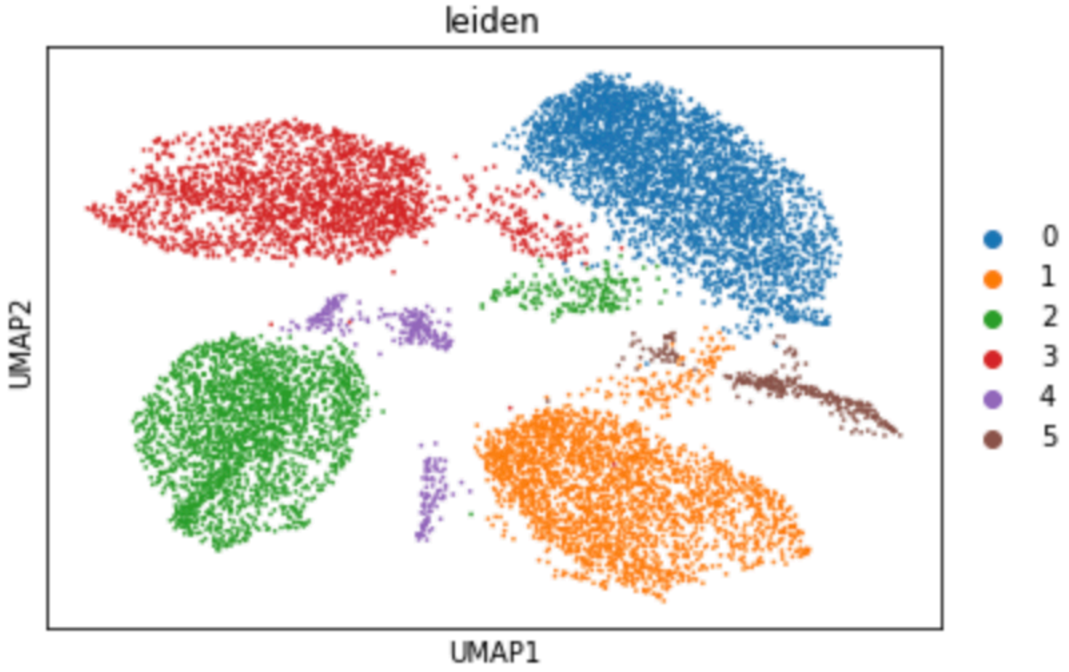


B

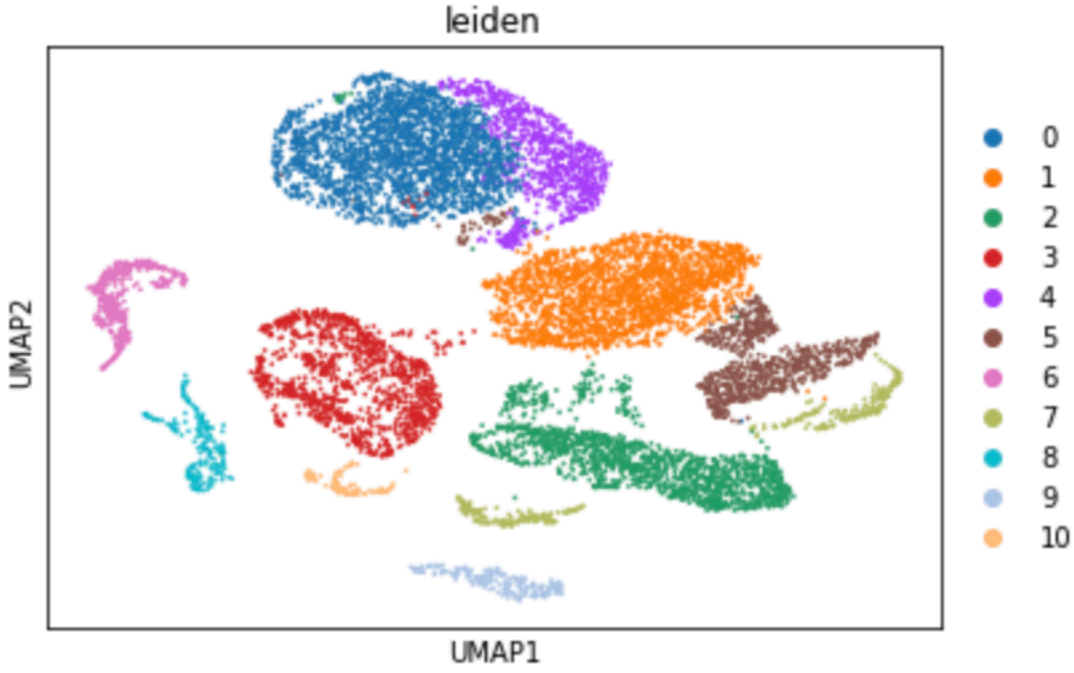


C

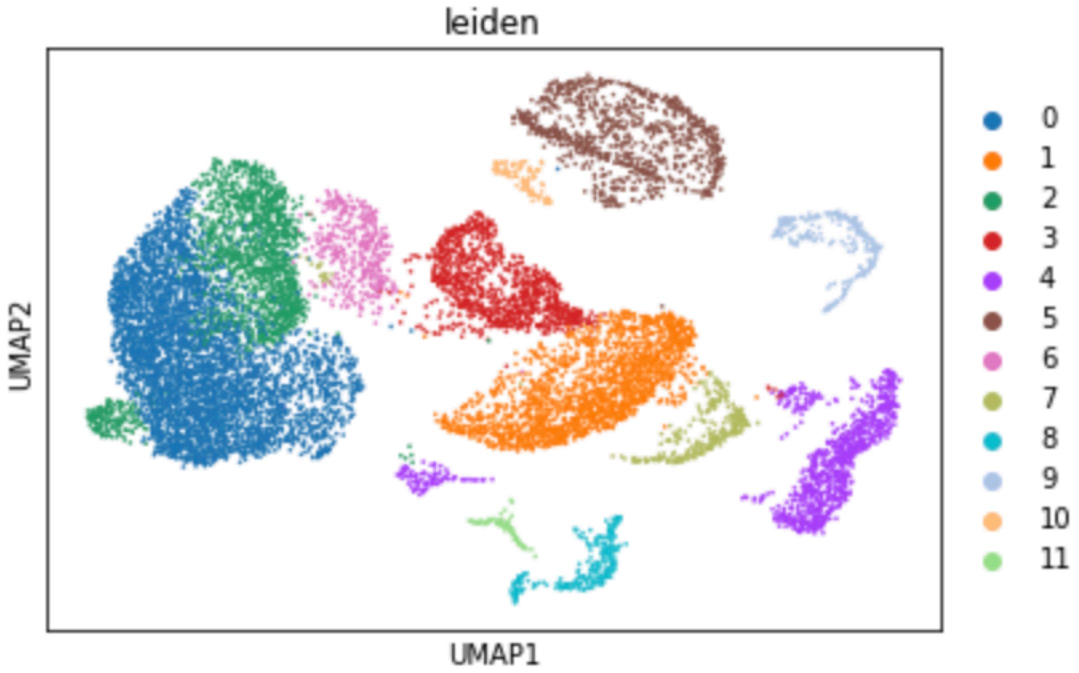
leiden
UMAP2
UMAP1
0
1
2
3
4
5
6
7
8
9
10
11

D

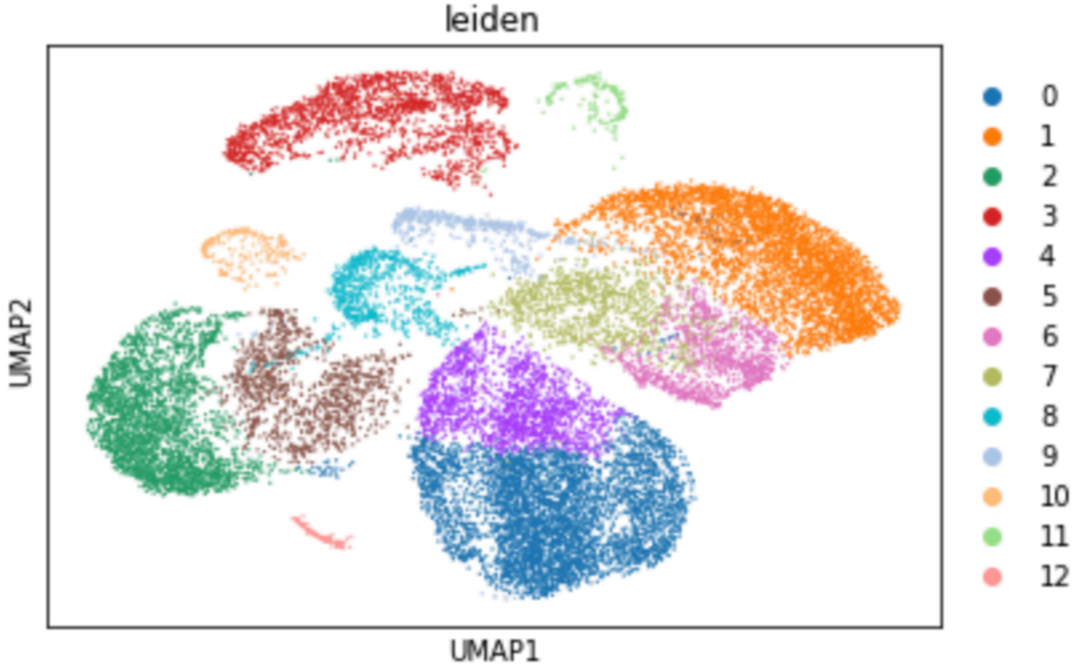
leiden
UMAP2
UMAP1
0
1
2
3
4
5
6
7
8
9
10
11
12

E

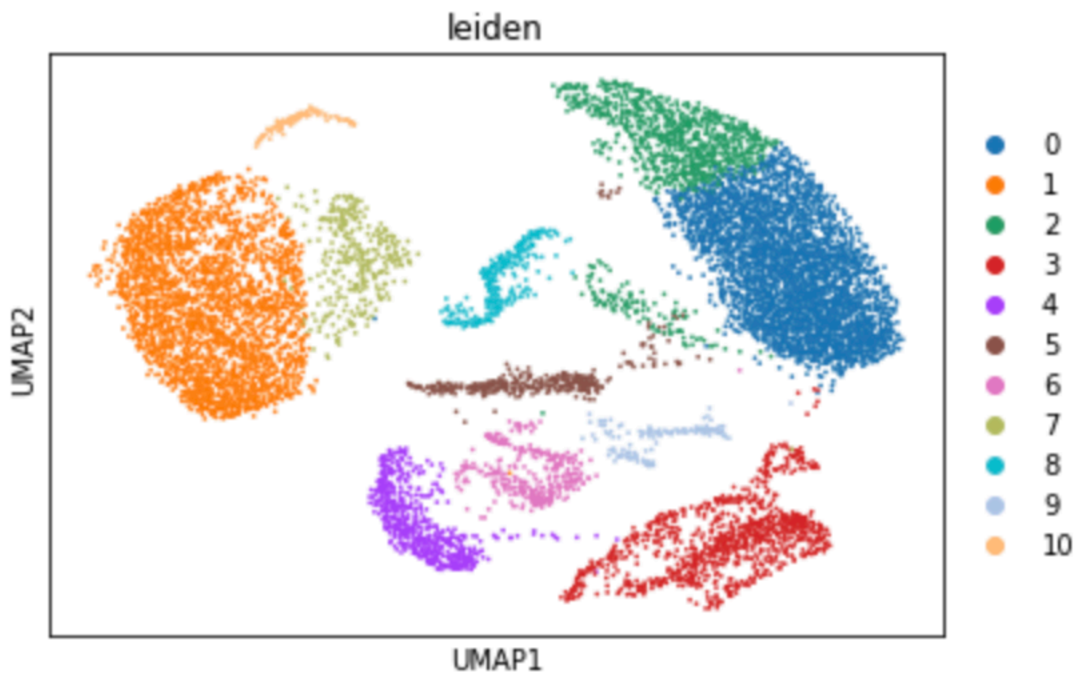


F

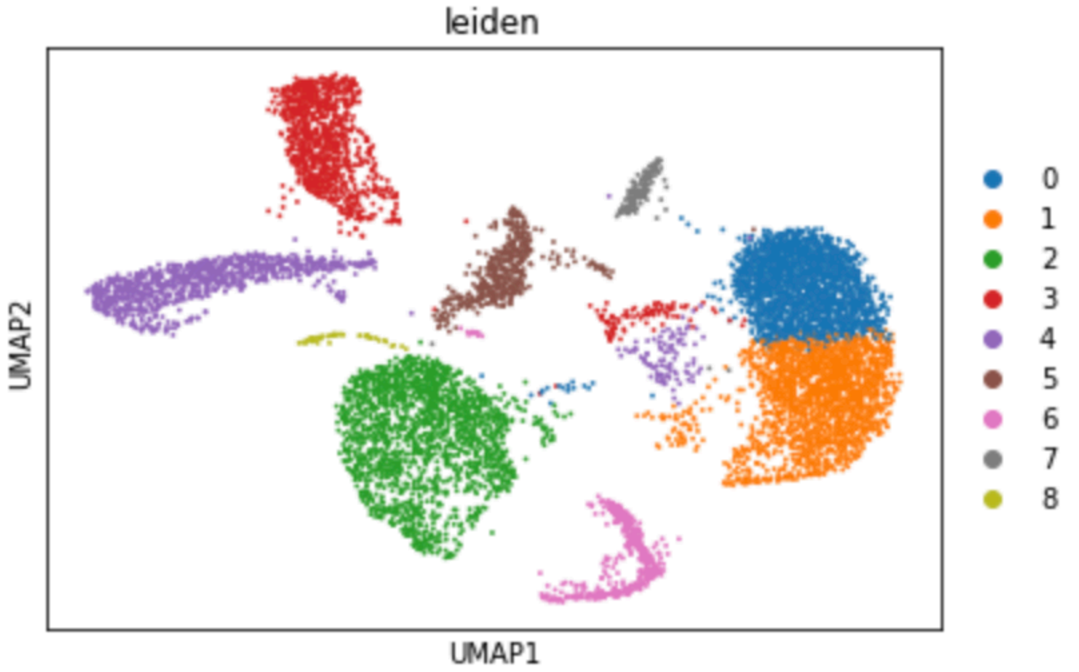


Legend: PCA based umap reduction visualization graph of PCA leiden method generated clusters. (A, B, C): control1, control2, control3, AD1, AD2, AD3.

Fig2:
A
1

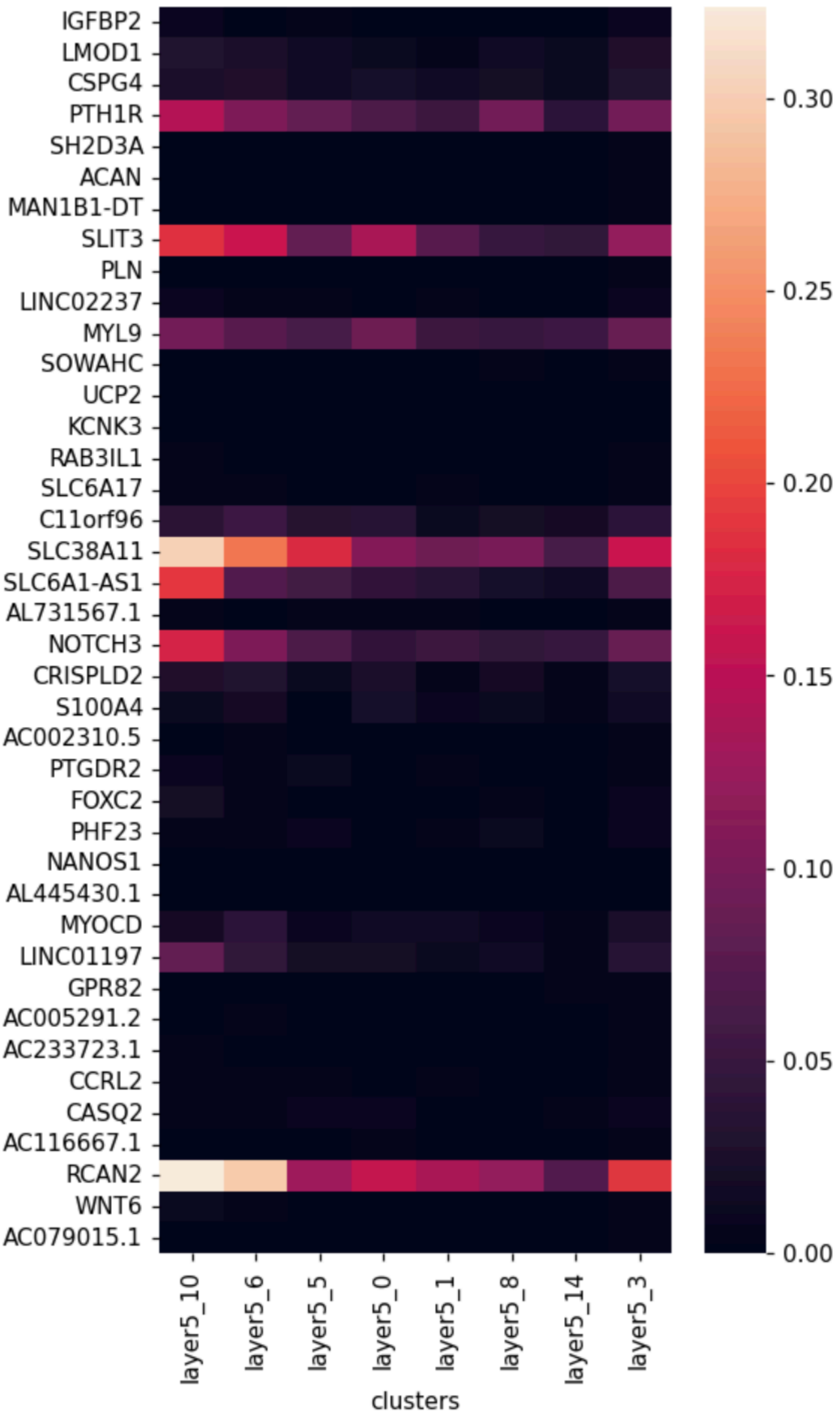
IGFBP2
LMOD1
CSPG4
PTH1R
SH2D3A
ACAN
MAN1B1-DT
SLIT3
PLN
LINC02237
MYL9
SOWAHC
UCP2
KCNK3
RAB3IL1
SLC6A17
C11orf96
SLC38A11
SLC6A1-AS1
AL731567.1
NOTCH3
CRISPLD2
S100A4
AC002310.5
PTGDR2
FOXC2
PHF23
NANOS1
AL445430.1
MYOCD
LINC01197
GPR82
AC005291.2
AC233723.1
CCRL2
CASQ2
AC116667.1
RCAN2
WNT6
AC079015.1
layer5_10
layer5_6
layer5_5
layer5_0
layer5_1
layer5_8
layer5_14
layer5_3
clusters
0.30
0.25
0.20
0.15
0.10
0.05
0.00

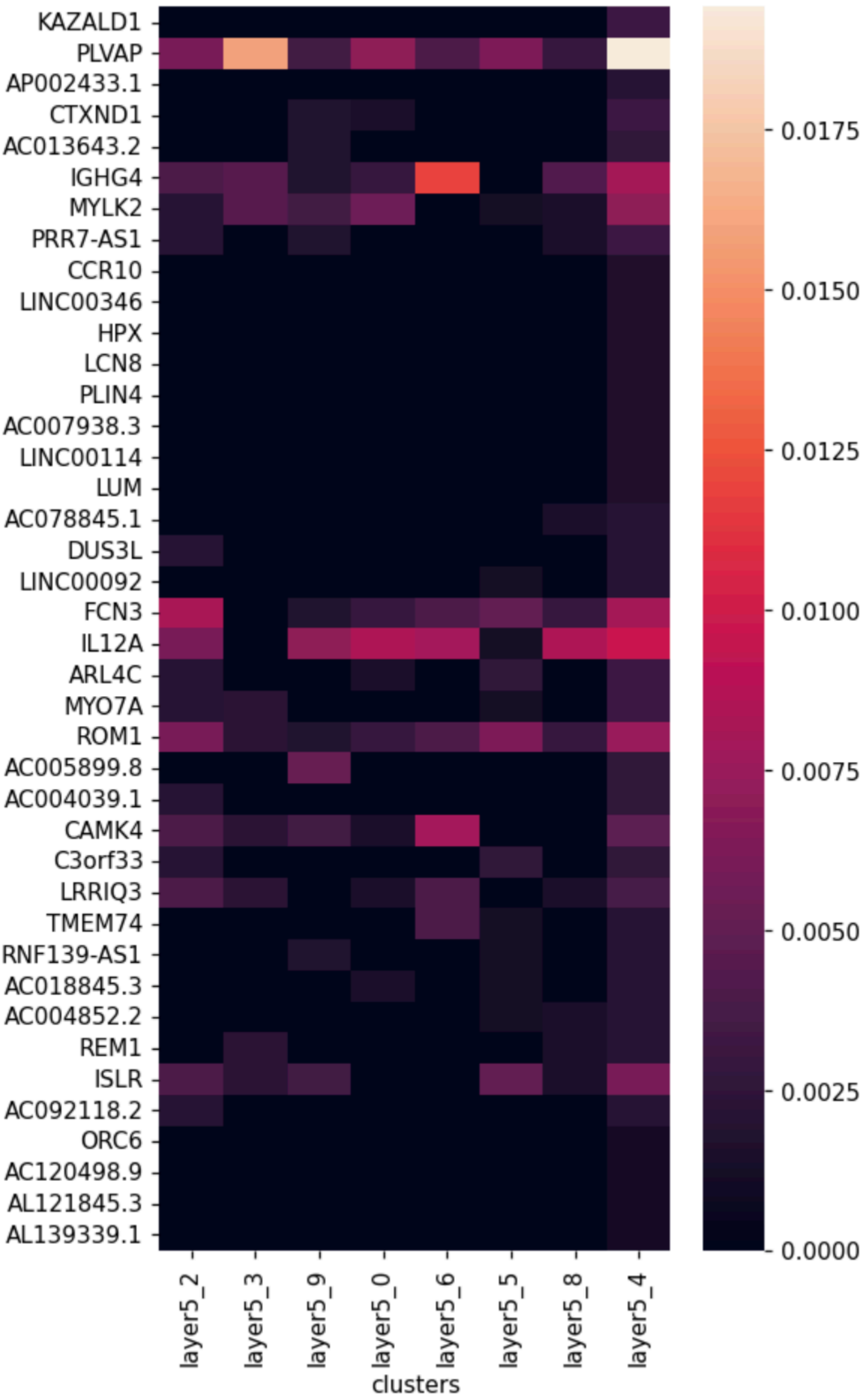

KAZALD1
PLVAP
AP002433.1
CTXND1
AC013643.2
IGHG4
MYLK2
PRR7-AS1
CCR10
LINC00346
HPX
LCN8
PLIN4
AC007938.3
LINC00114
LUM
AC078845.1
DUS3L
LINC00092
FCN3
IL12A
ARL4C
MYO7A
ROM1
AC005899.8
AC004039.1
CAMK4
C3orf33
LRRIQ3
TMEM74
RNF139-AS1
AC018845.3
AC004852.2
REM1
ISLR
AC092118.2
ORC6
AC120498.9
AL121845.3
AL139339.1
layer5_2
layer5_3
layer5_9
layer5_0
layer5_6
layer5_5
layer5_8
layer5_4
clusters
0.0175
0.0150
0.0125
0.0100
0.0075
0.0050
0.0025
0.0000

B
1

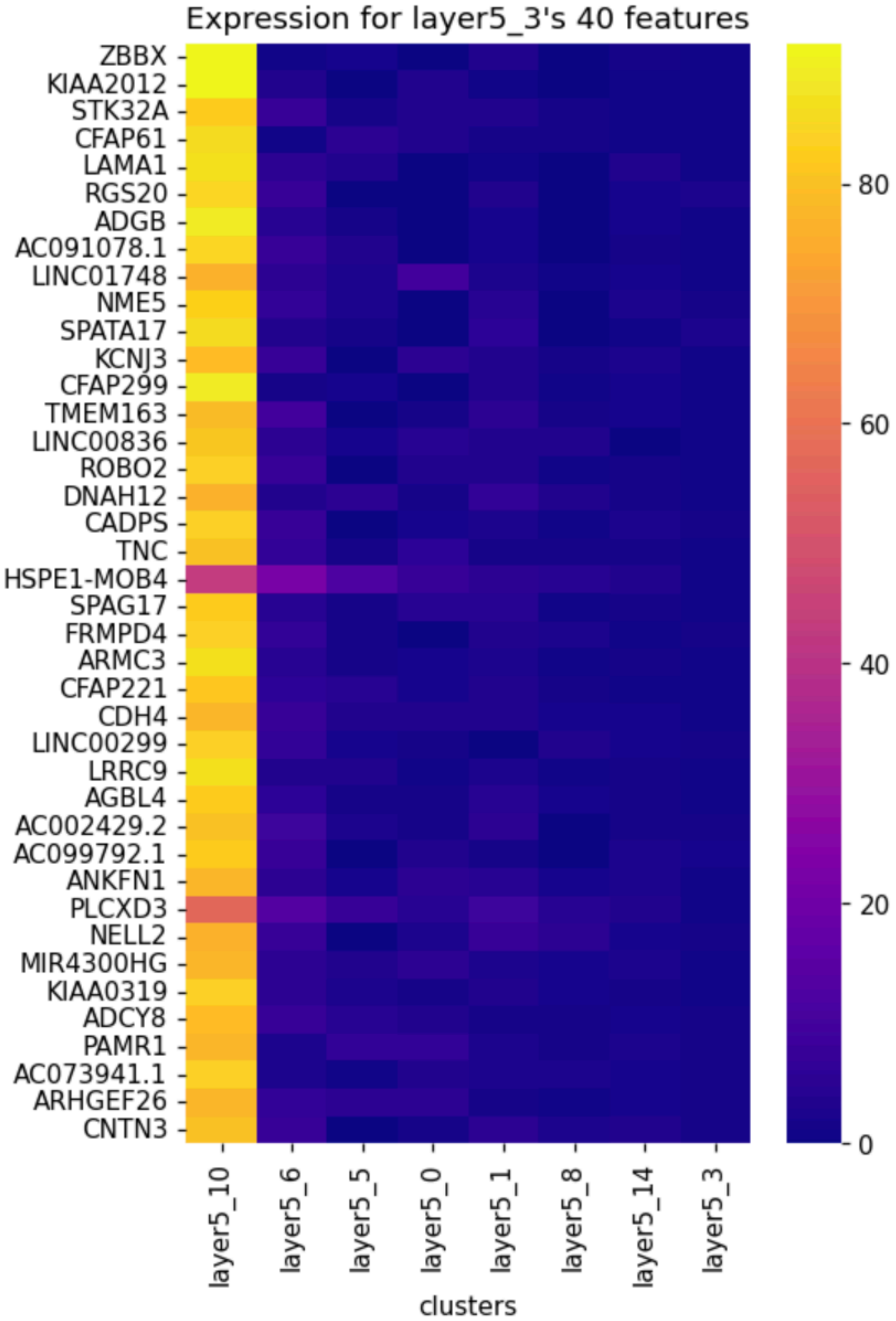
Expression for layer5_3's 40 features
ZBBX
KIAA2012
STK32A
CFAP61
LAMA1
RGS20
ADGB
AC091078.1
LINC01748
NME5
SPATA17
KCNJ3
CFAP299
TMEM163
LINC00836
ROBO2
DNAH12
CADPS
TNC
HSPE1-MOB4
SPAG17
FRMPD4
ARMC3
CFAP221
CDH4
LINC00299
LRRC9
AGBL4
AC002429.2
AC099792.1
ANKFN1
PLCXD3
NELL2
MIR4300HG
KIAA0319
ADCY8
PAMR1
AC073941.1
ARHGEF26
CNTN3
layer5_10
layer5_6
layer5_5
layer5_0
layer5_1
layer5_8
layer5_14
layer5_3
clusters
80
60
40
20
0

2

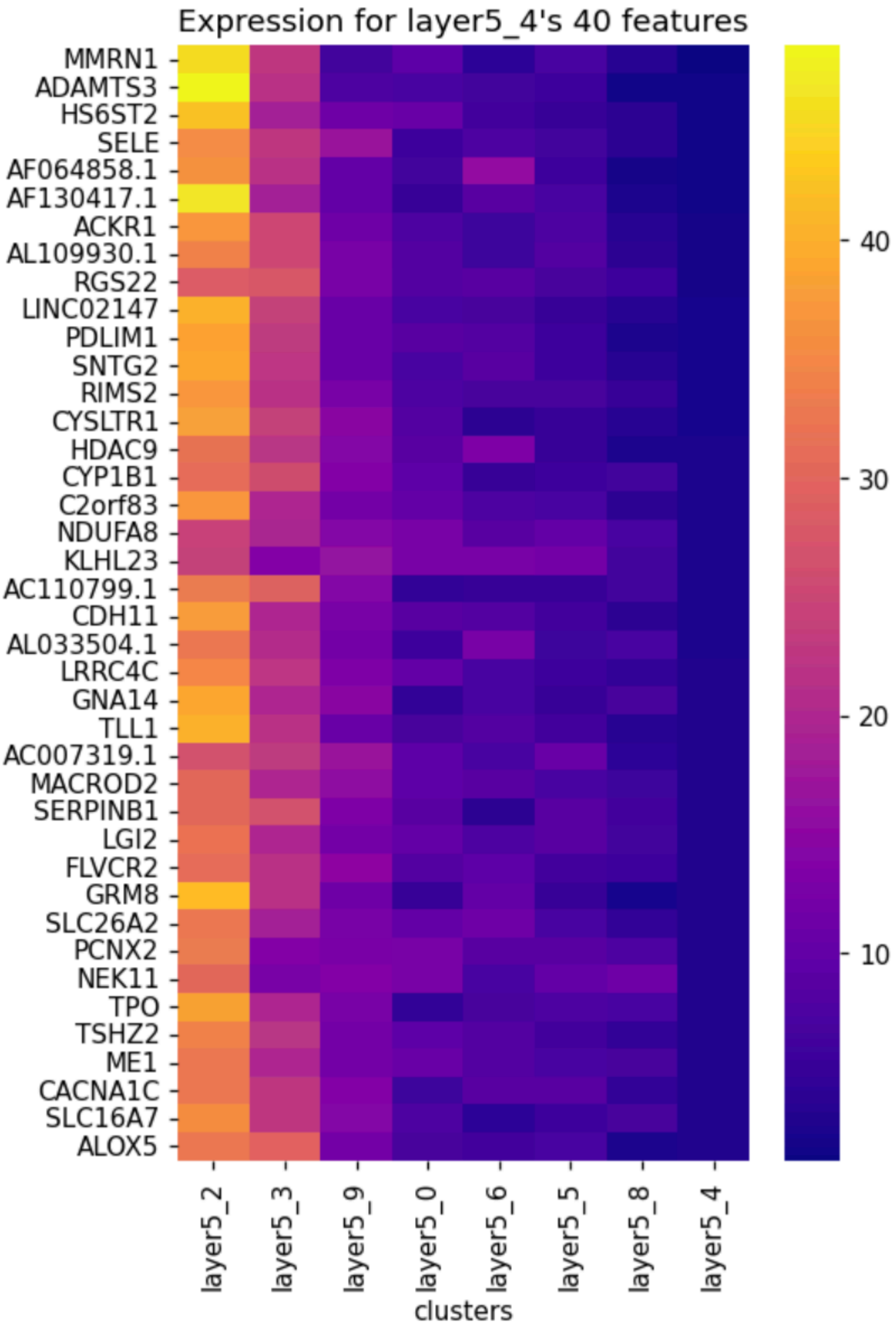
Expression for layer5_4's 40 features
MMRN1
ADAMTS3
HS6ST2
SELE
AF064858.1
AF130417.1
ACKR1
AL109930.1
RGS22
LINC02147
PDLIM1
SNTG2
RIMS2
CYSLTR1
HDAC9
CYP1B1
C2orf83
NDUFA8
KLHL23
AC110799.1
CDH11
AL033504.1
LRRC4C
GNA14
TLL1
AC007319.1
MACROD2
SERPINB1
LGI2
FLVCR2
GRM8
SLC26A2
PCNX2
NEK11
TPO
TSHZ2
ME1
CACNA1C
SLC16A7
ALOX5
layer5_2
layer5_3
layer5_9
layer5_0
layer5_6
layer5_5
layer5_8
layer5_4
clusters
40
30
20
10

Legend: heatmaps of top 40 marker genes of the latest aging stage. (A, B): Scanpy rank_gene_group function result, BW high variable function result. (1, 2): AD1 sample, control3 sample.

Fig3

A

1

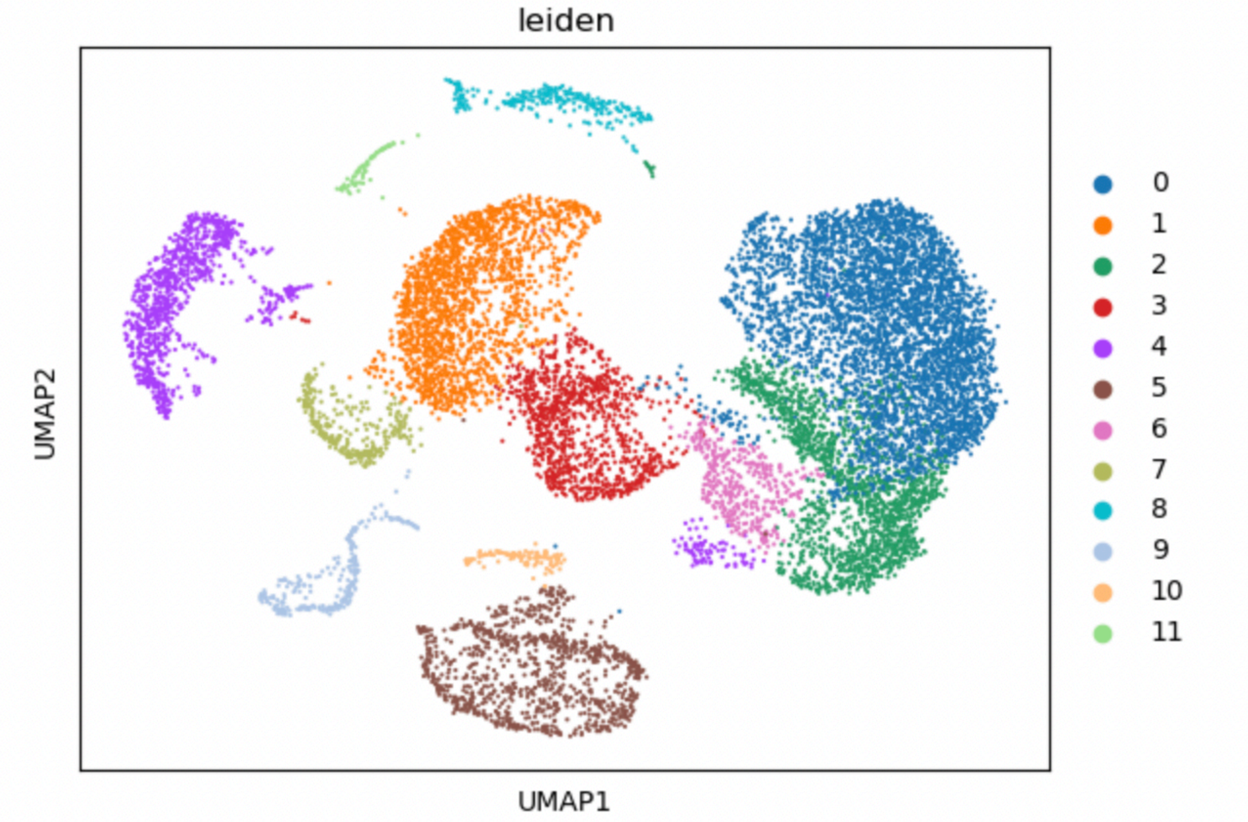


2

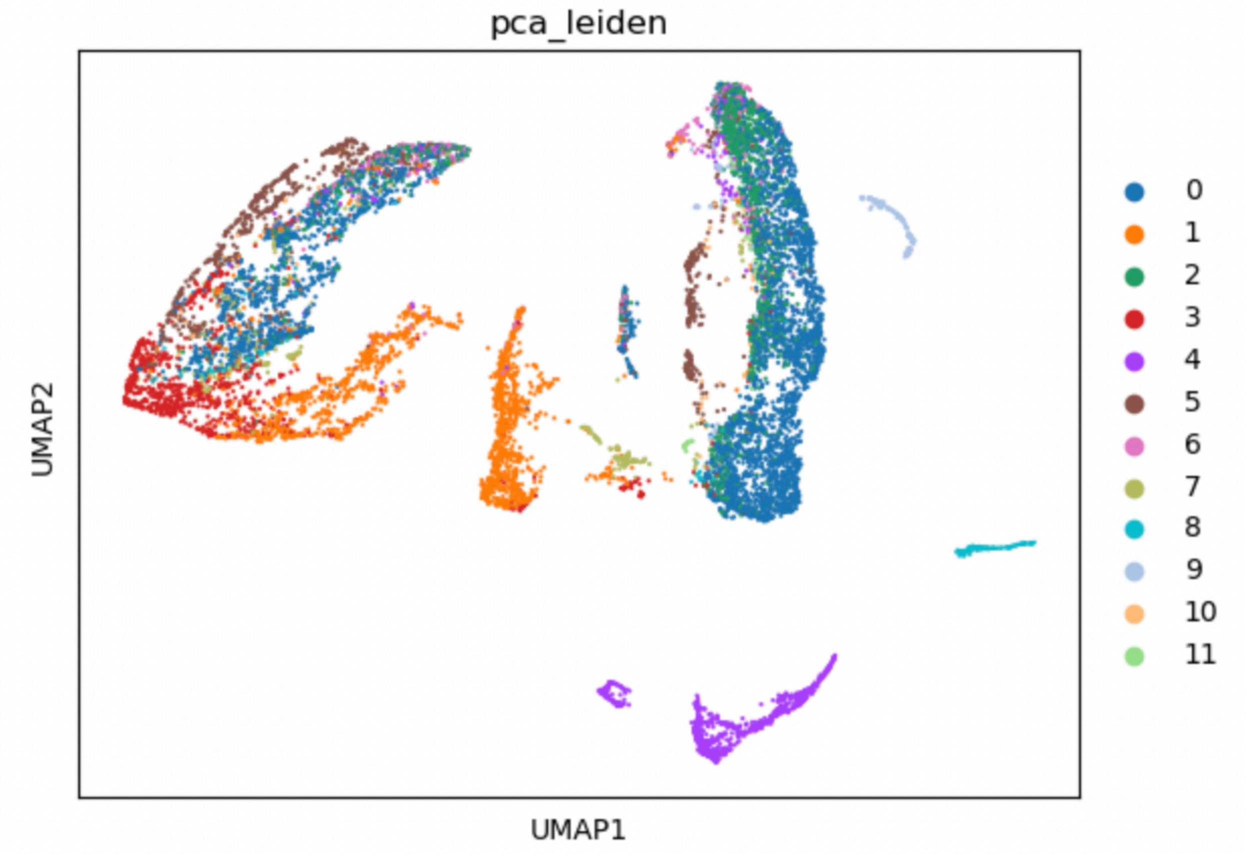
pca_leiden
UMAP2
UMAP1
0
1
2
3
4
5
6
7
8
9
10
11

3

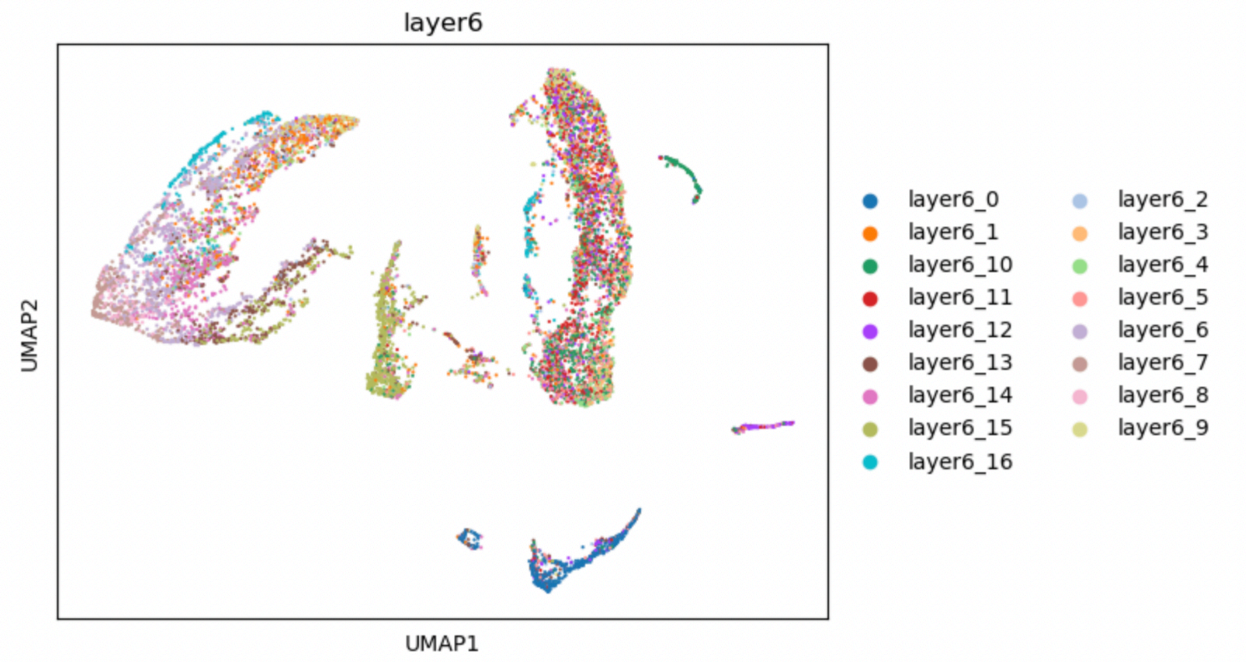
layer6
UMAP2
UMAP1
layer6_0
layer6_1
layer6_10
layer6_11
layer6_12
layer6_13
layer6_14
layer6_15
layer6_16
layer6_2
layer6_3
layer6_4
layer6_5
layer6_6
layer6_7
layer6_8
layer6_9

4

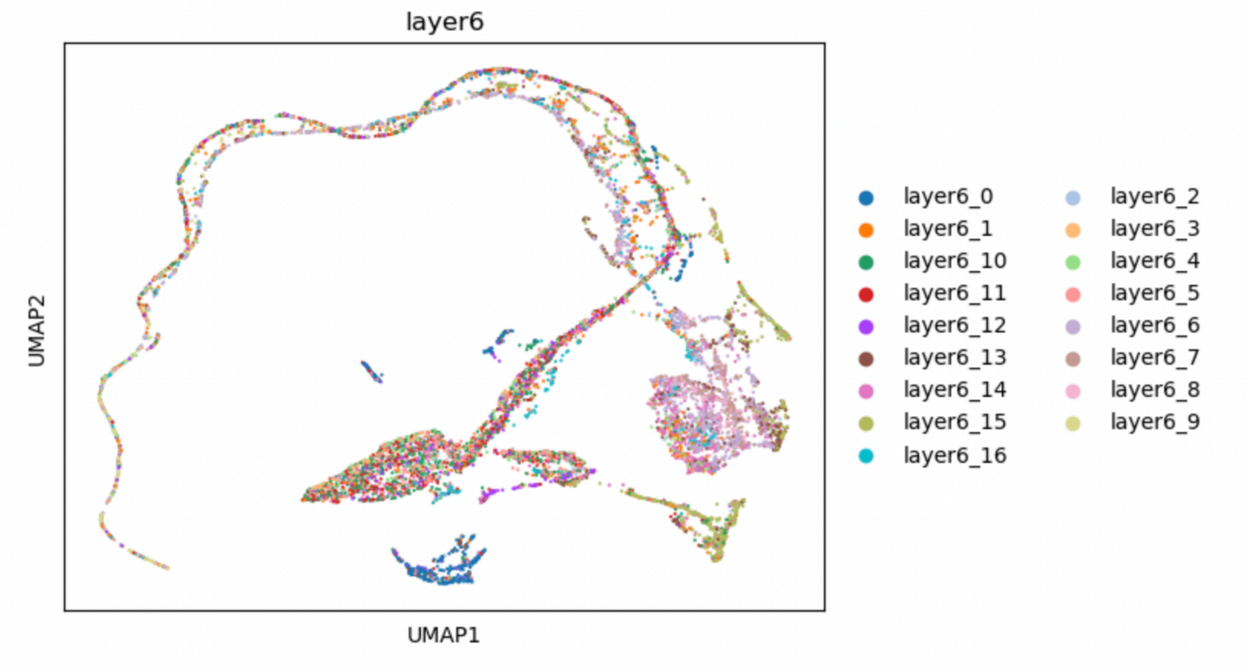
layer6
UMAP2
UMAP1
layer6_0
layer6_1
layer6_10
layer6_11
layer6_12
layer6_13
layer6_14
layer6_15
layer6_16
layer6_2
layer6_3
layer6_4
layer6_5
layer6_6
layer6_7
layer6_8
layer6_9

5

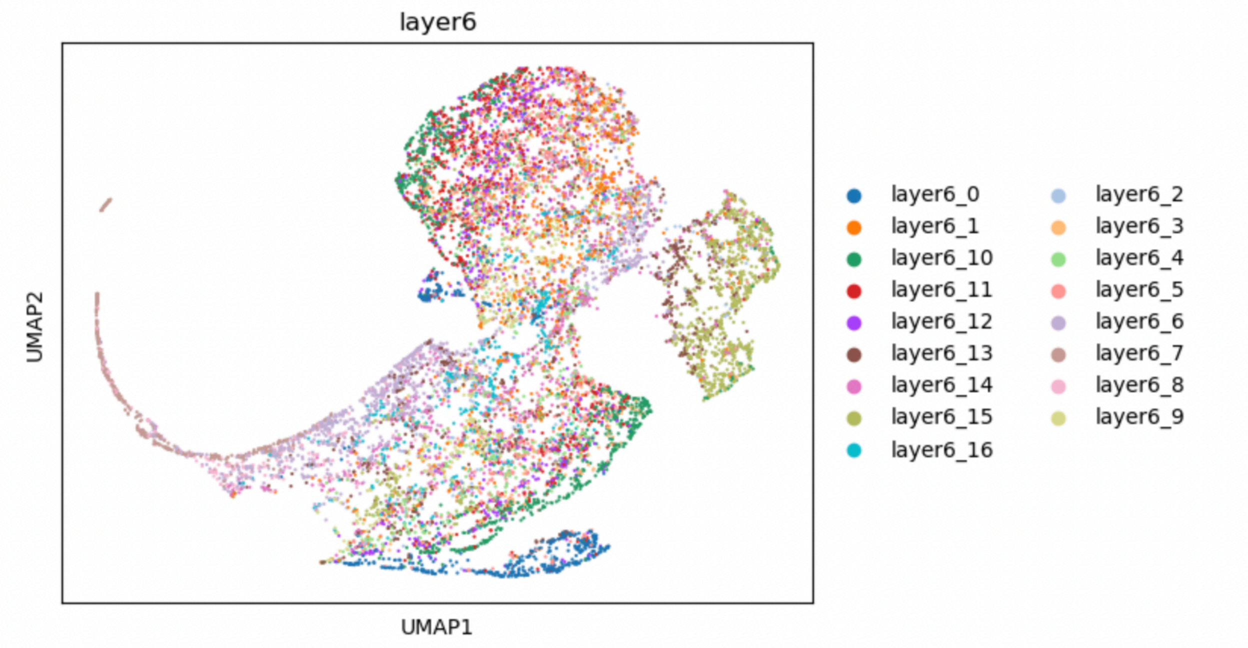
layer6
UMAP2
UMAP1
layer6_0
layer6_1
layer6_10
layer6_11
layer6_12
layer6_13
layer6_14
layer6_15
layer6_16
layer6_2
layer6_3
layer6_4
layer6_5
layer6_6
layer6_7
layer6_8
layer6_9

B

1

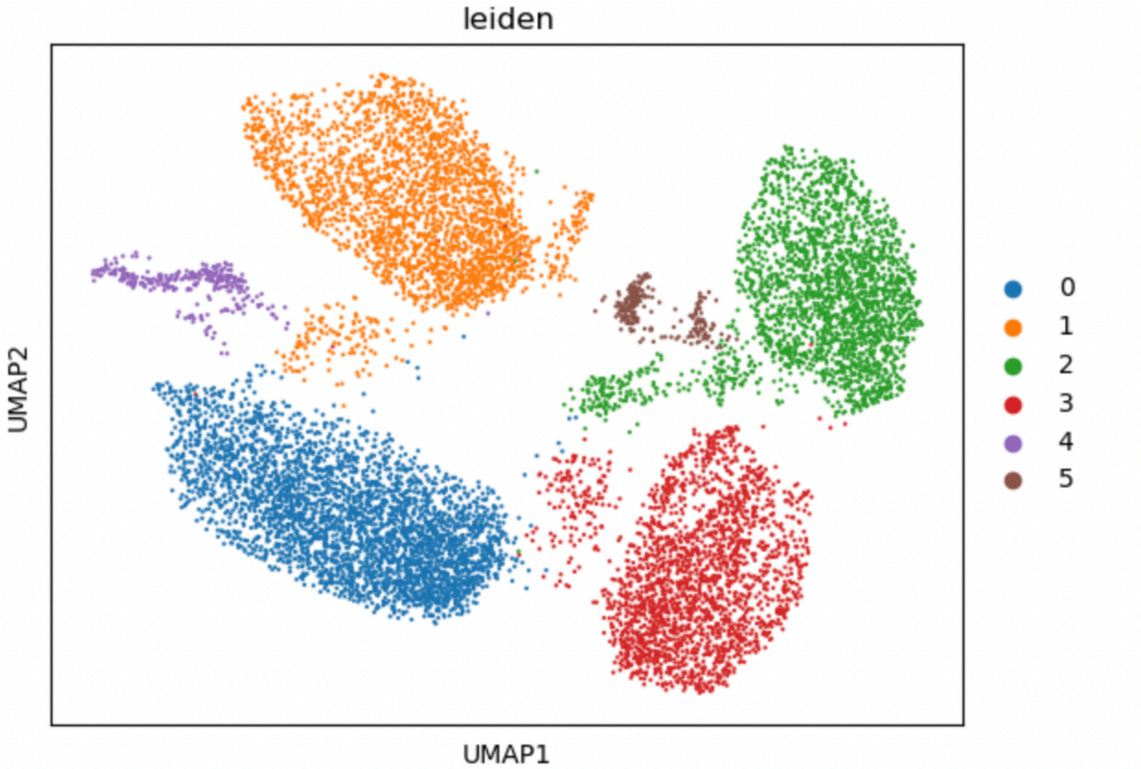


2

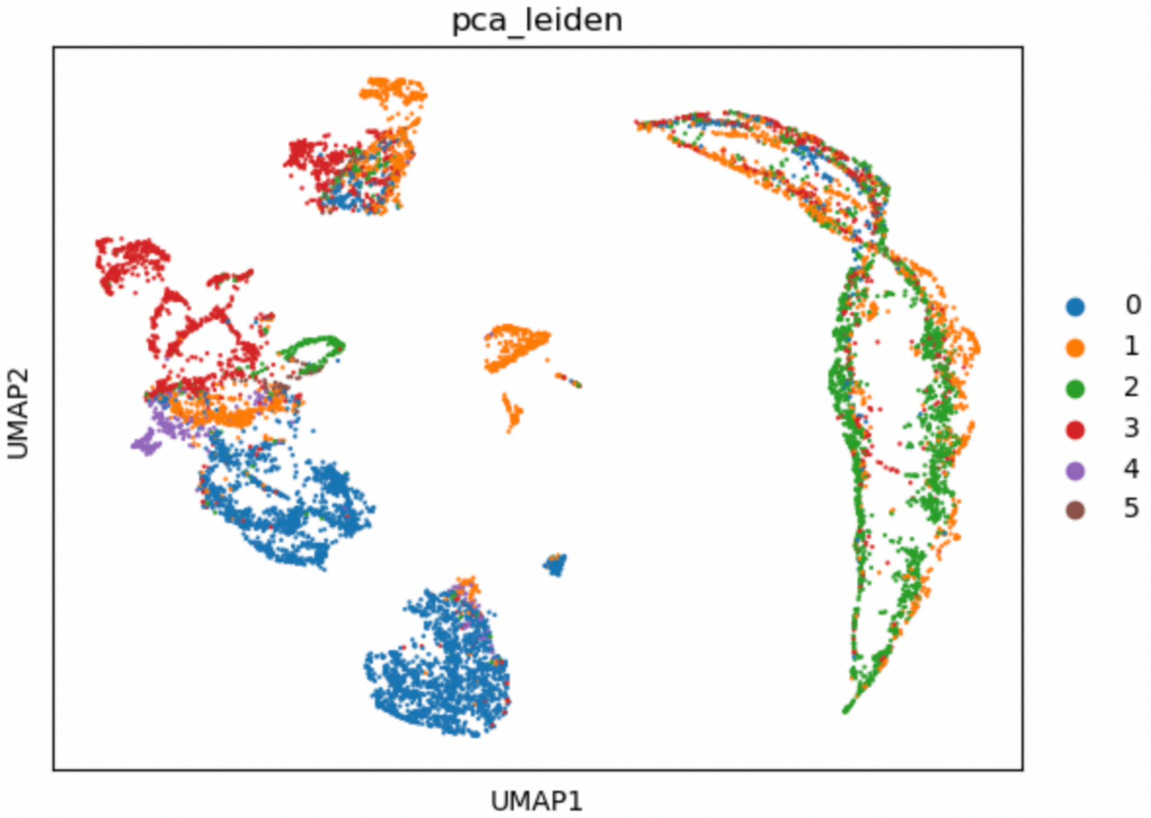


3

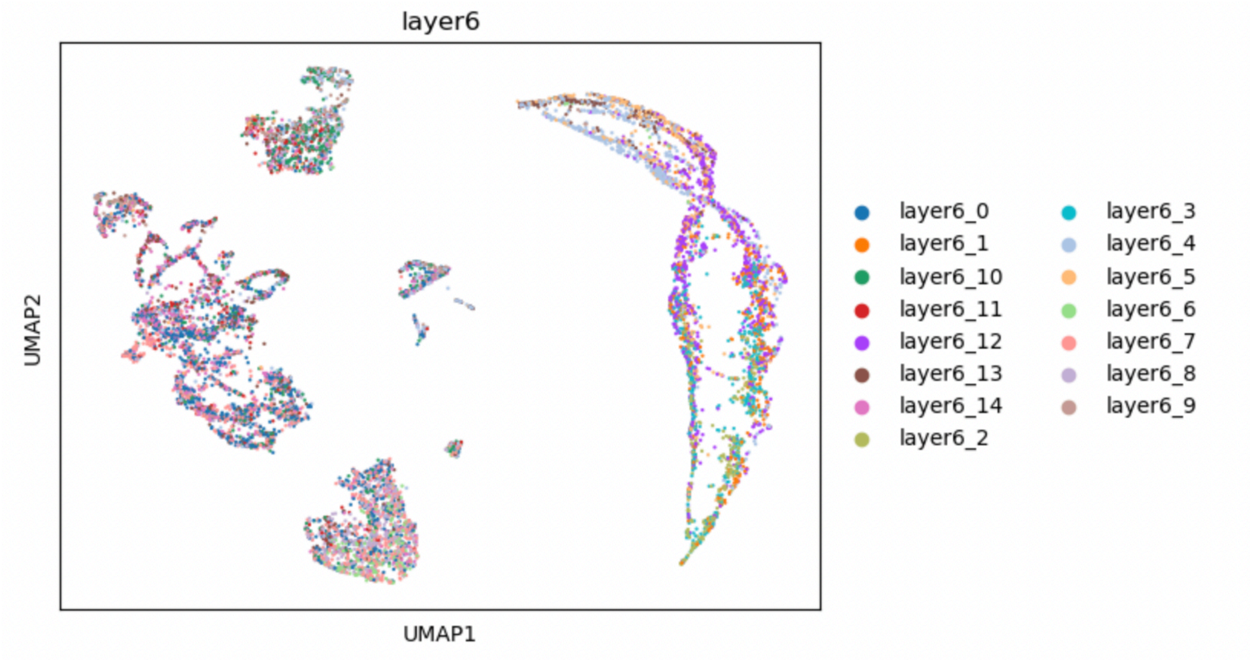
layer6
UMAP2
UMAP1
layer6_0
layer6_1
layer6_10
layer6_11
layer6_12
layer6_13
layer6_14
layer6_2
layer6_3
layer6_4
layer6_5
layer6_6
layer6_7
layer6_8
layer6_9

4

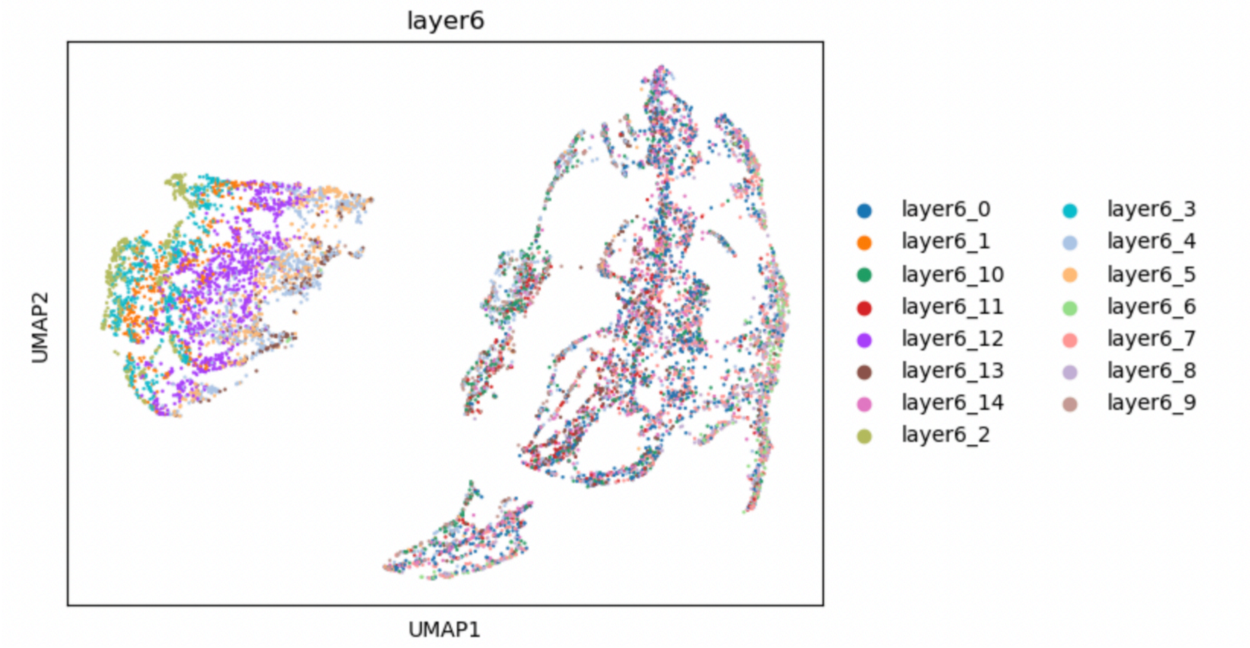
layer6
UMAP2
UMAP1
layer6_0
layer6_1
layer6_10
layer6_11
layer6_12
layer6_13
layer6_14
layer6_2
layer6_3
layer6_4
layer6_5
layer6_6
layer6_7
layer6_8
layer6_9

5

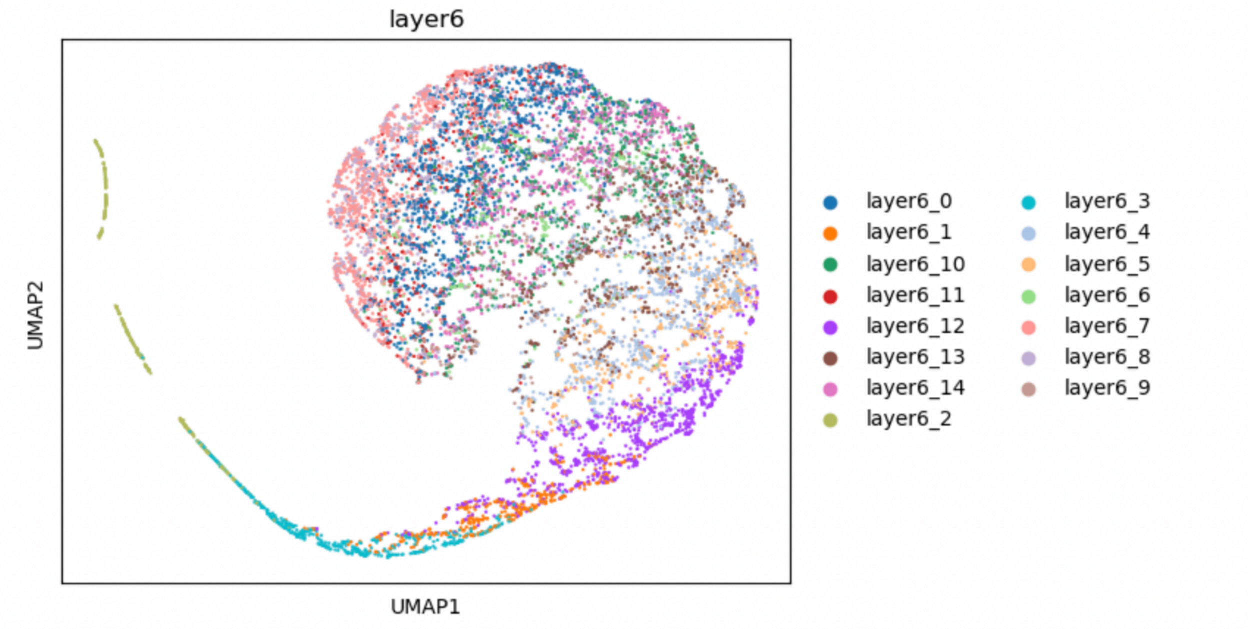
layer6
UMAP2
UMAP1
layer6_0
layer6_1
layer6_10
layer6_11
layer6_12
layer6_13
layer6_14
layer6_2
layer6_3
layer6_4
layer6_5
layer6_6
layer6_7
layer6_8
layer6_9

C

1

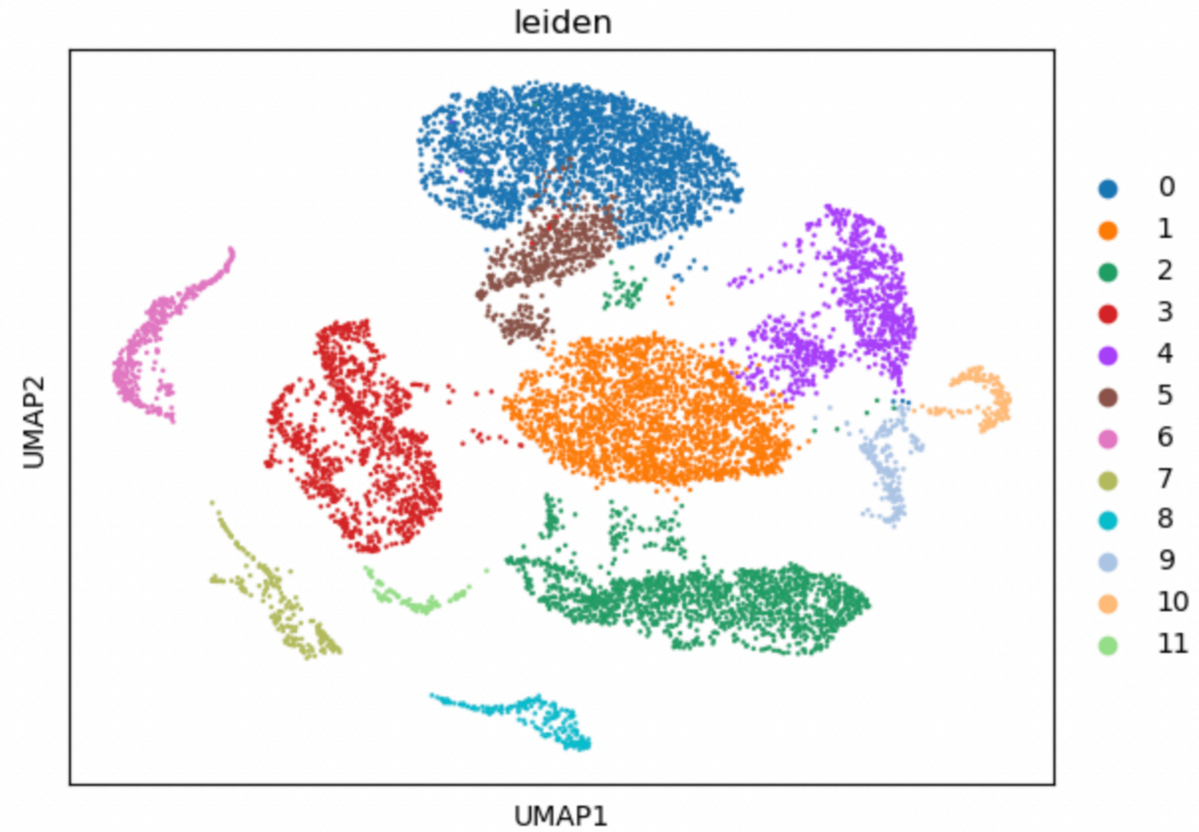
leiden
UMAP2
UMAP1
0
1
2
3
4
5
6
7
8
9
10
11

2

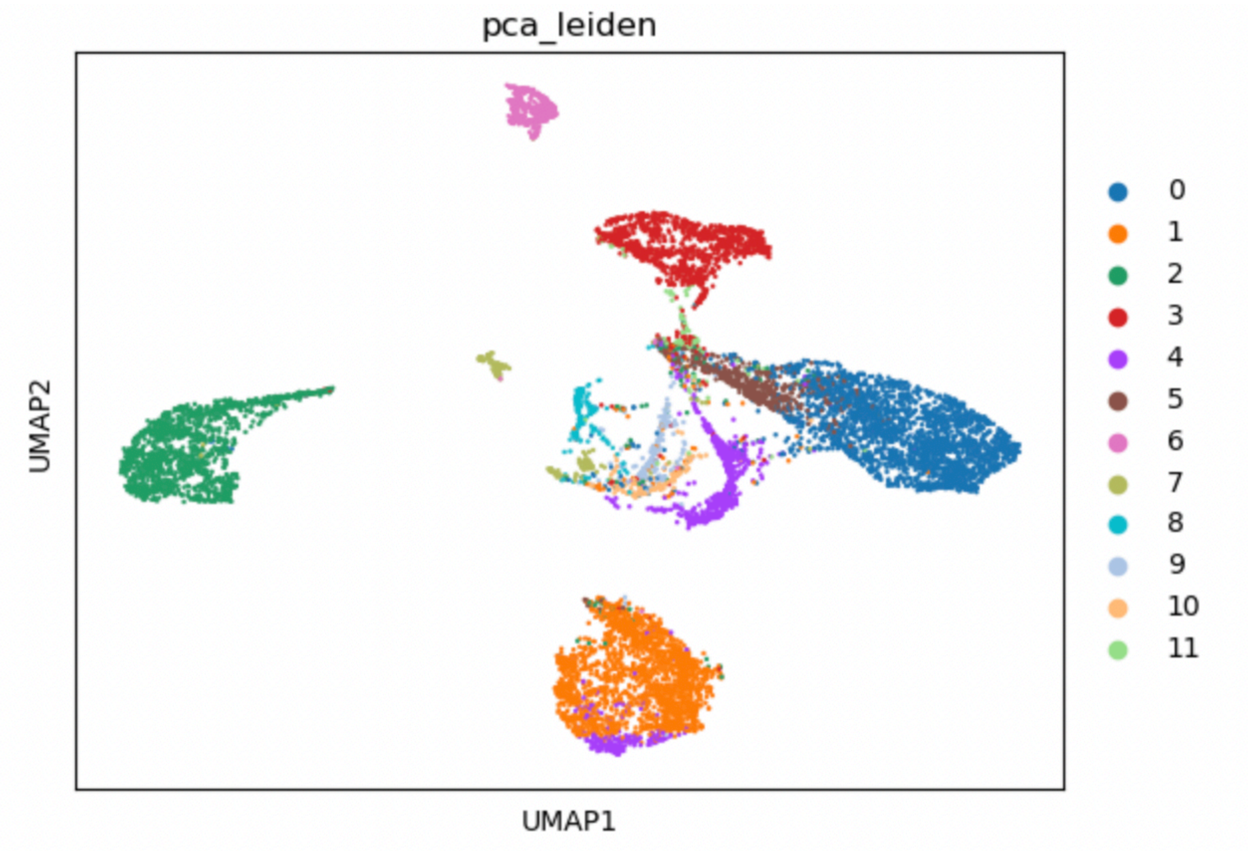
pca_leiden
UMAP2
UMAP1
0
1
2
3
4
5
6
7
8
9
10
11

3

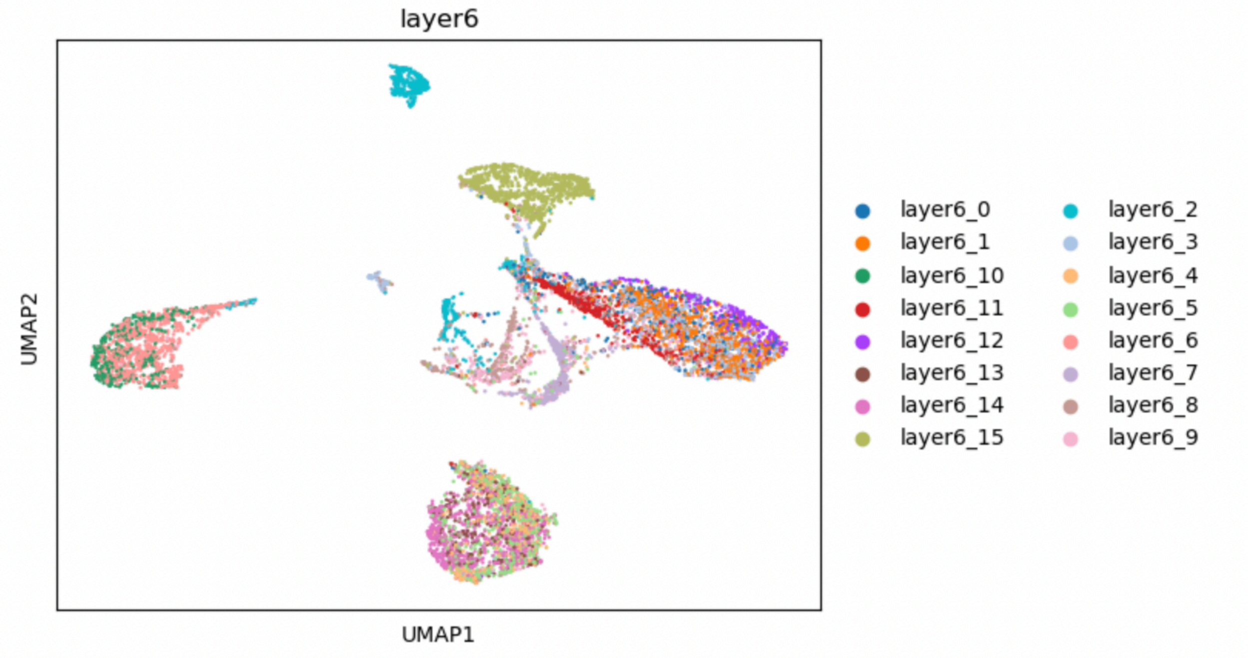
layer6
UMAP2
UMAP1
layer6_0
layer6_1
layer6_10
layer6_11
layer6_12
layer6_13
layer6_14
layer6_15
layer6_2
layer6_3
layer6_4
layer6_5
layer6_6
layer6_7
layer6_8
layer6_9

4

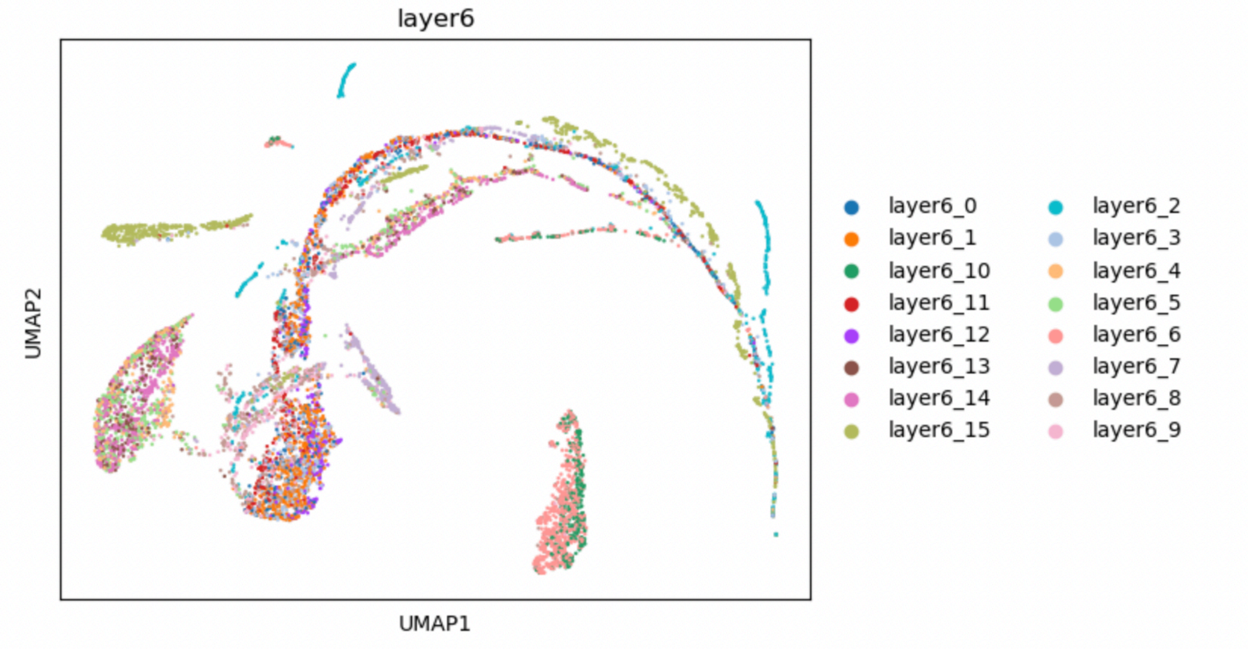
layer6
UMAP2
UMAP1
layer6_0
layer6_1
layer6_10
layer6_11
layer6_12
layer6_13
layer6_14
layer6_15
layer6_2
layer6_3
layer6_4
layer6_5
layer6_6
layer6_7
layer6_8
layer6_9

5

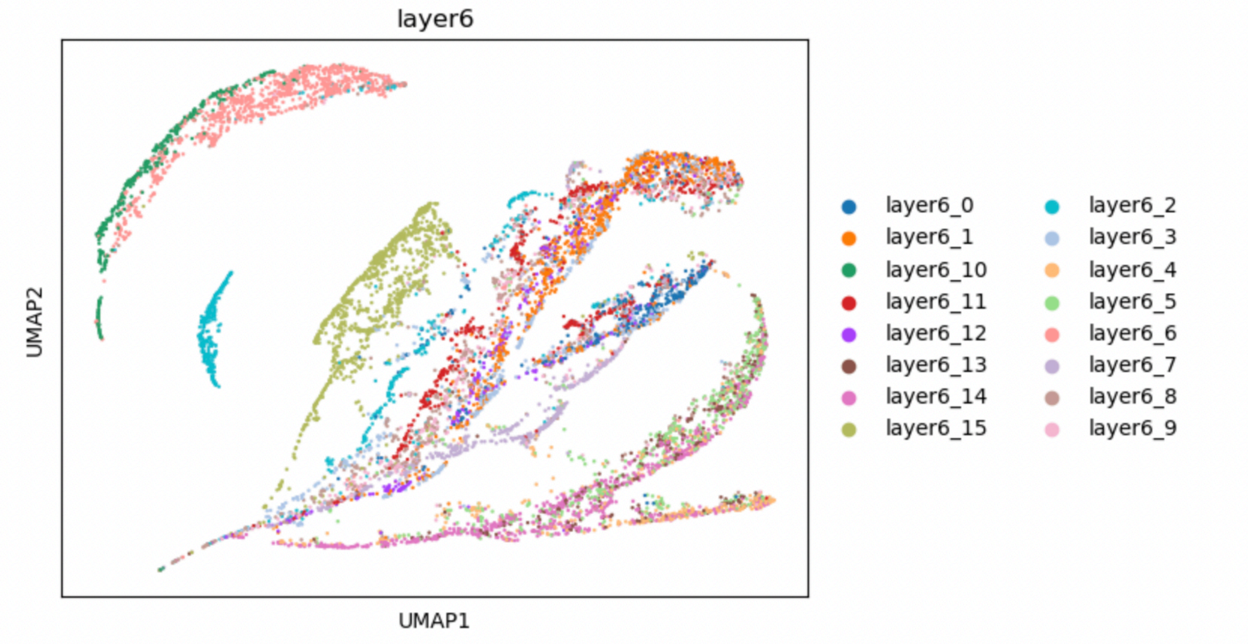
layer6
UMAP2
UMAP1
layer6_0
layer6_1
layer6_10
layer6_11
layer6_12
layer6_13
layer6_14
layer6_15
layer6_2
layer6_3
layer6_4
layer6_5
layer6_6
layer6_7
layer6_8
layer6_9

D

1

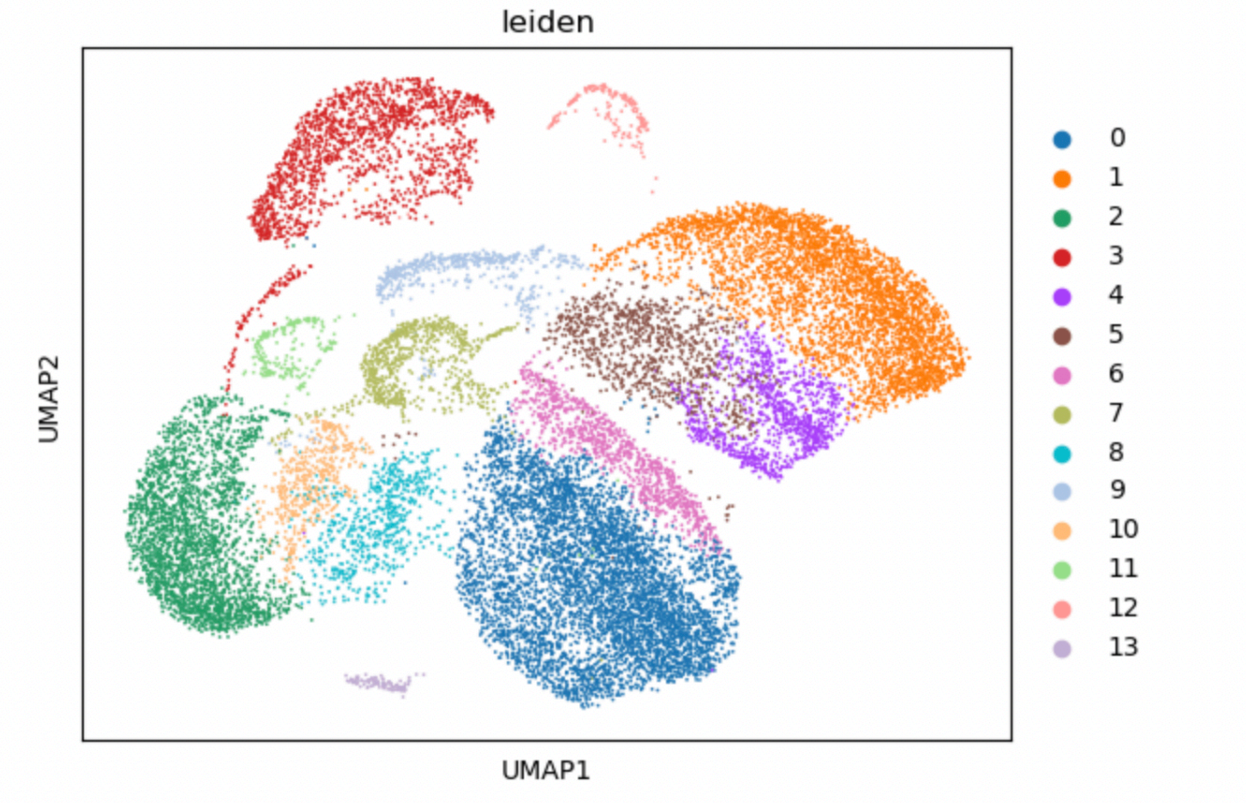
leiden
UMAP2
UMAP1
0
1
2
3
4
5
6
7
8
9
10
11
12
13

2

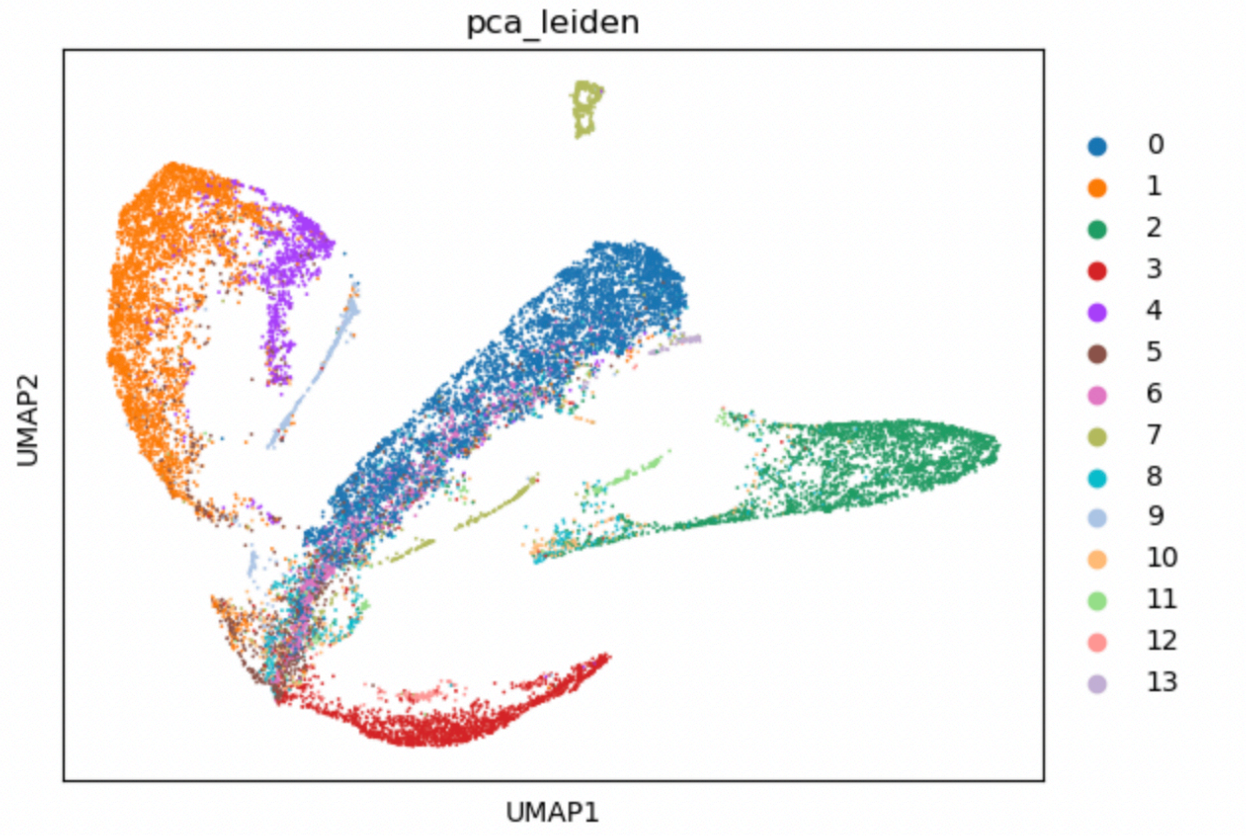
pca_leiden
UMAP2
UMAP1
0
1
2
3
4
5
6
7
8
9
10
11
12
13

3

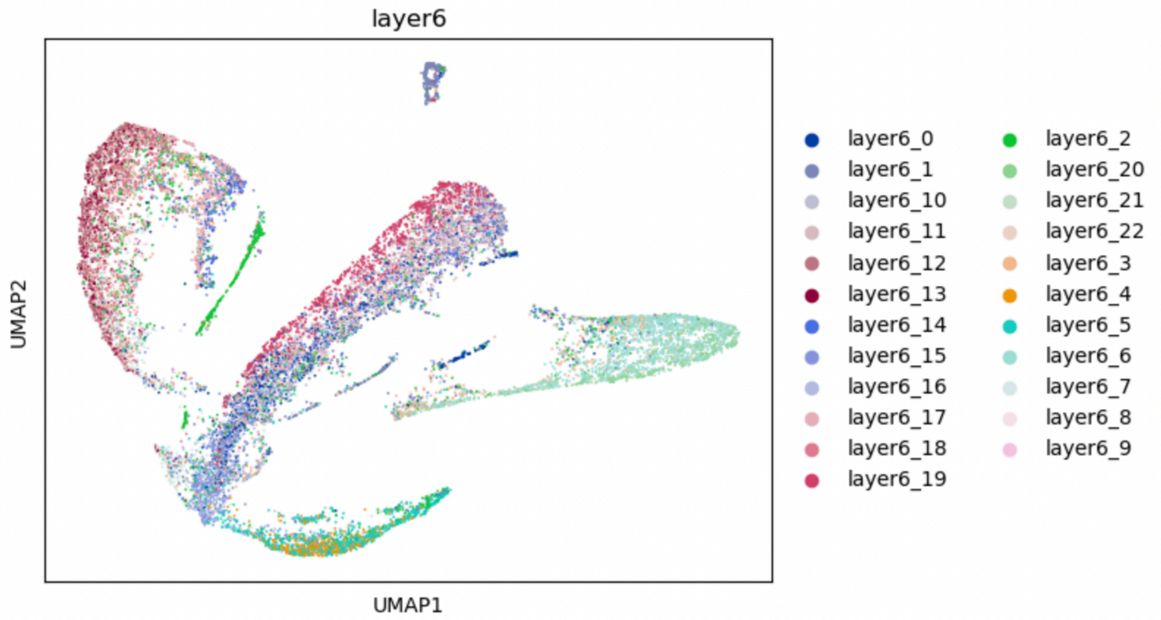
layer6
UMAP2
UMAP1
layer6_0
layer6_1
layer6_10
layer6_11
layer6_12
layer6_13
layer6_14
layer6_15
layer6_16
layer6_17
layer6_18
layer6_19
layer6_2
layer6_20
layer6_21
layer6_22
layer6_3
layer6_4
layer6_5
layer6_6
layer6_7
layer6_8
layer6_9

4

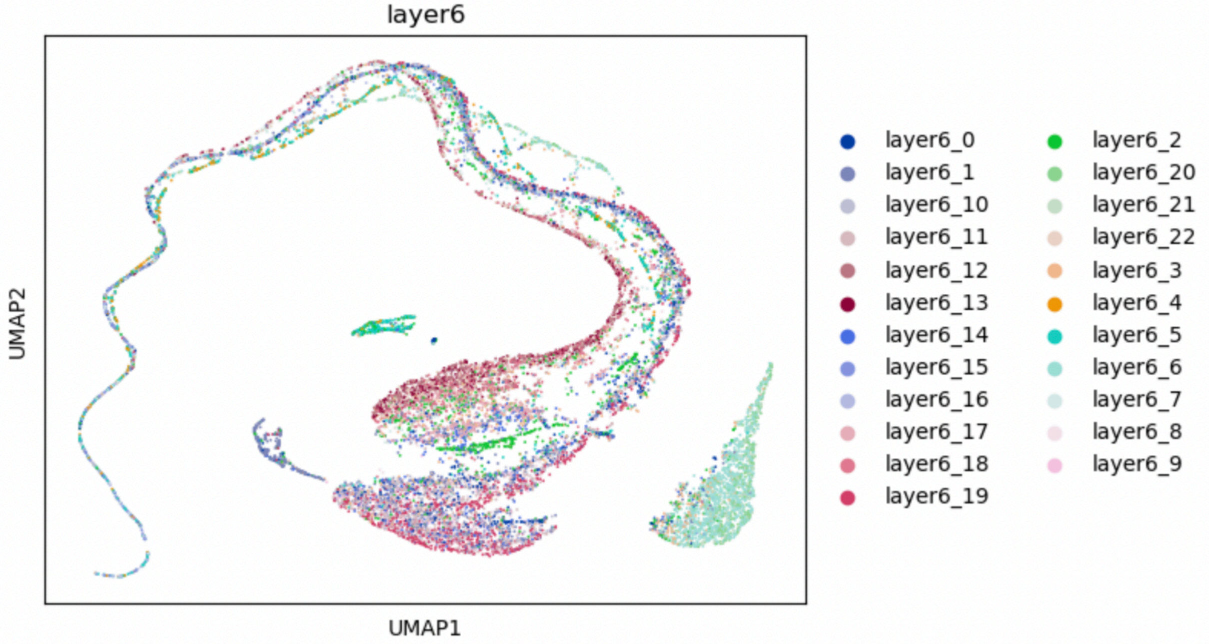
layer6
UMAP2
UMAP1
layer6_0
layer6_1
layer6_10
layer6_11
layer6_12
layer6_13
layer6_14
layer6_15
layer6_16
layer6_17
layer6_18
layer6_19
layer6_2
layer6_20
layer6_21
layer6_22
layer6_3
layer6_4
layer6_5
layer6_6
layer6_7
layer6_8
layer6_9

5

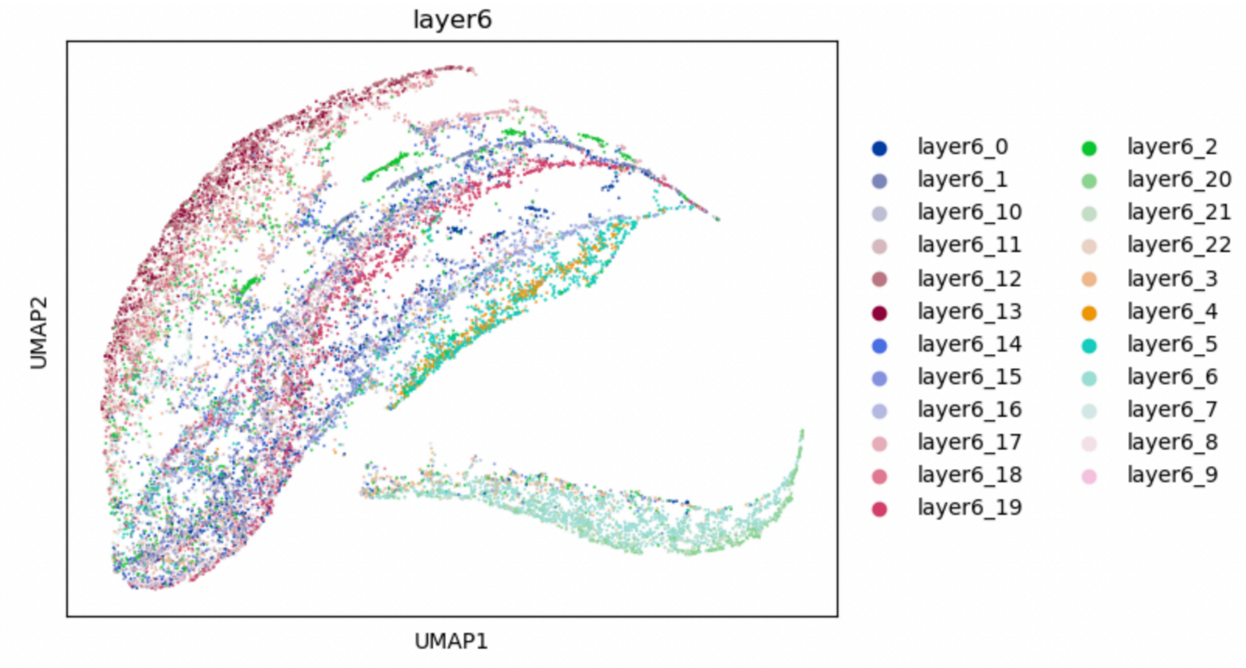
layer6
UMAP2
UMAP1
layer6_0
layer6_1
layer6_10
layer6_11
layer6_12
layer6_13
layer6_14
layer6_15
layer6_16
layer6_17
layer6_18
layer6_19
layer6_2
layer6_20
layer6_21
layer6_22
layer6_3
layer6_4
layer6_5
layer6_6
layer6_7
layer6_8
layer6_9

E

1

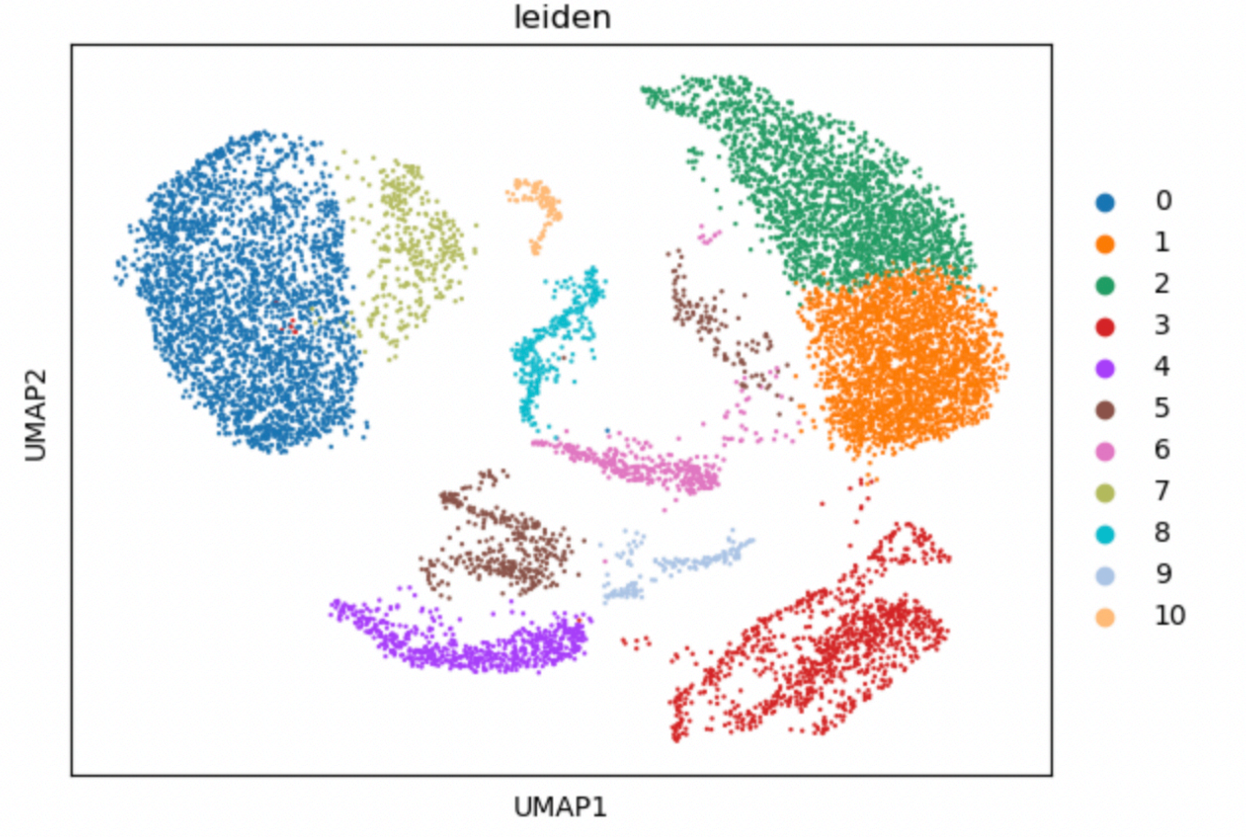
leiden
UMAP2
UMAP1
0
1
2
3
4
5
6
7
8
9
10

2

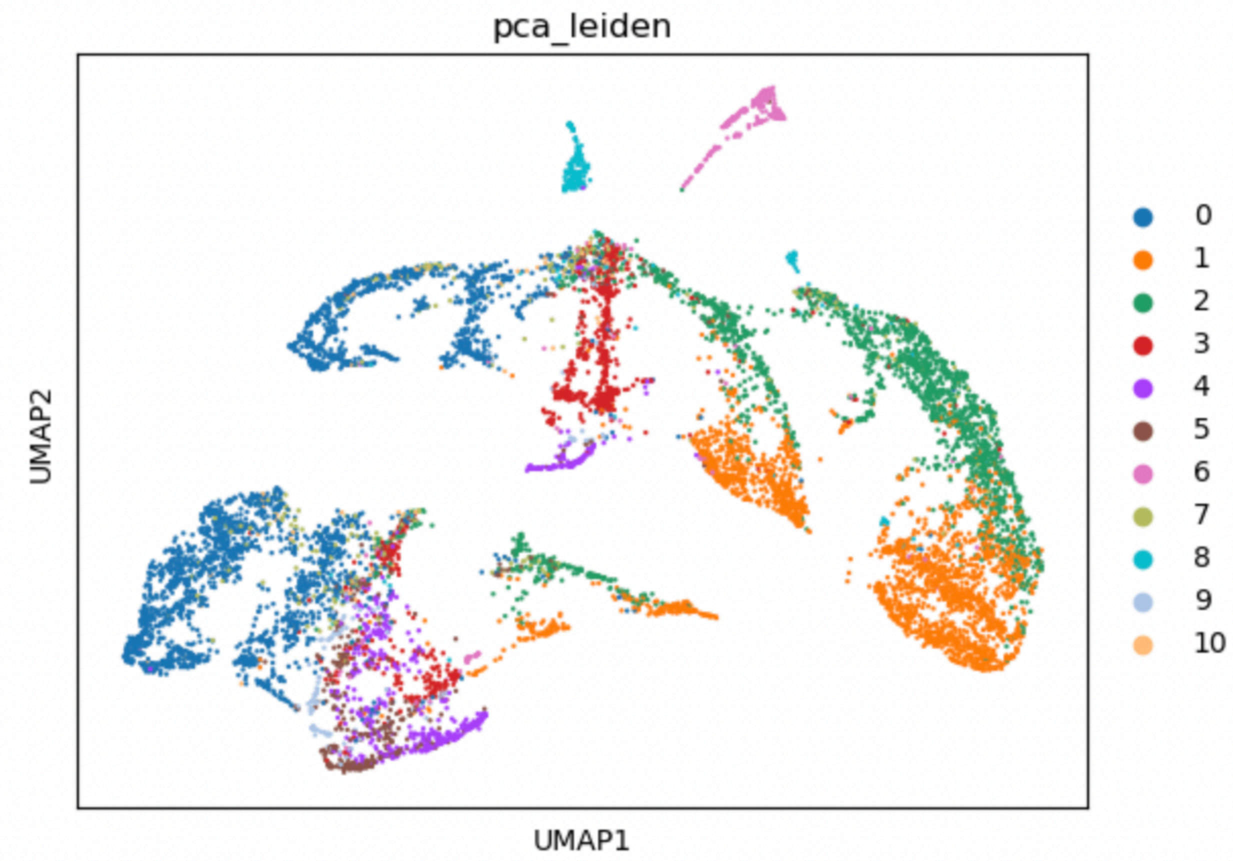
pca_leiden
UMAP2
UMAP1
0
1
2
3
4
5
6
7
8
9
10

3

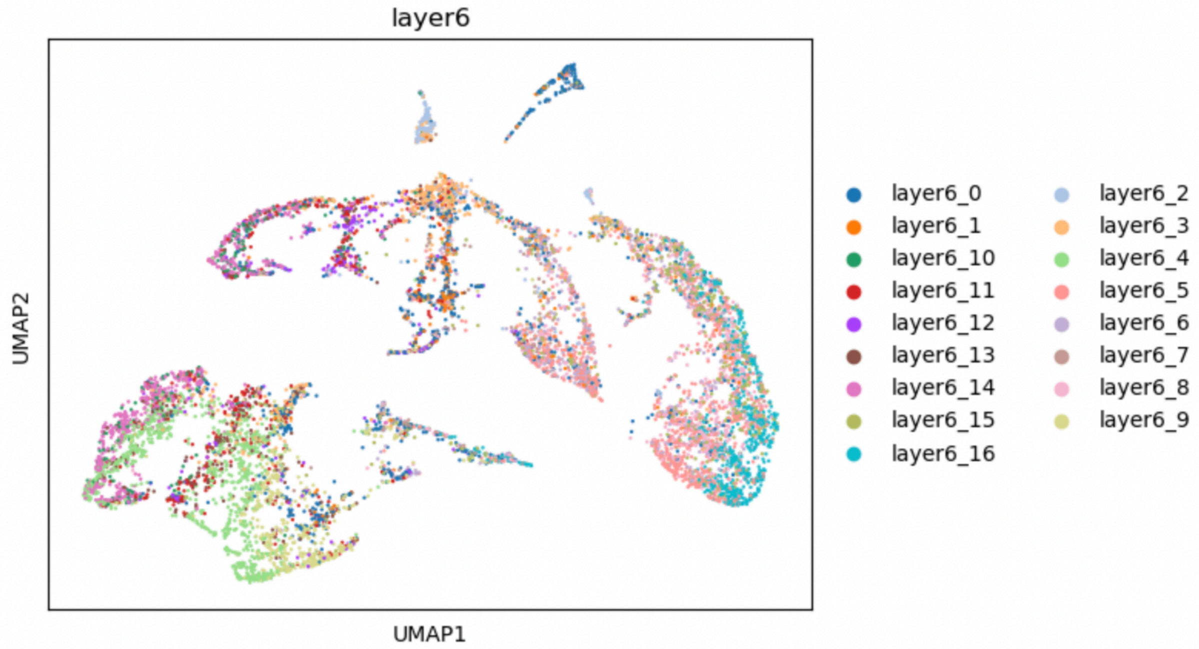
layer6
UMAP2
UMAP1
layer6_0
layer6_1
layer6_10
layer6_11
layer6_12
layer6_13
layer6_14
layer6_15
layer6_16
layer6_2
layer6_3
layer6_4
layer6_5
layer6_6
layer6_7
layer6_8
layer6_9

4

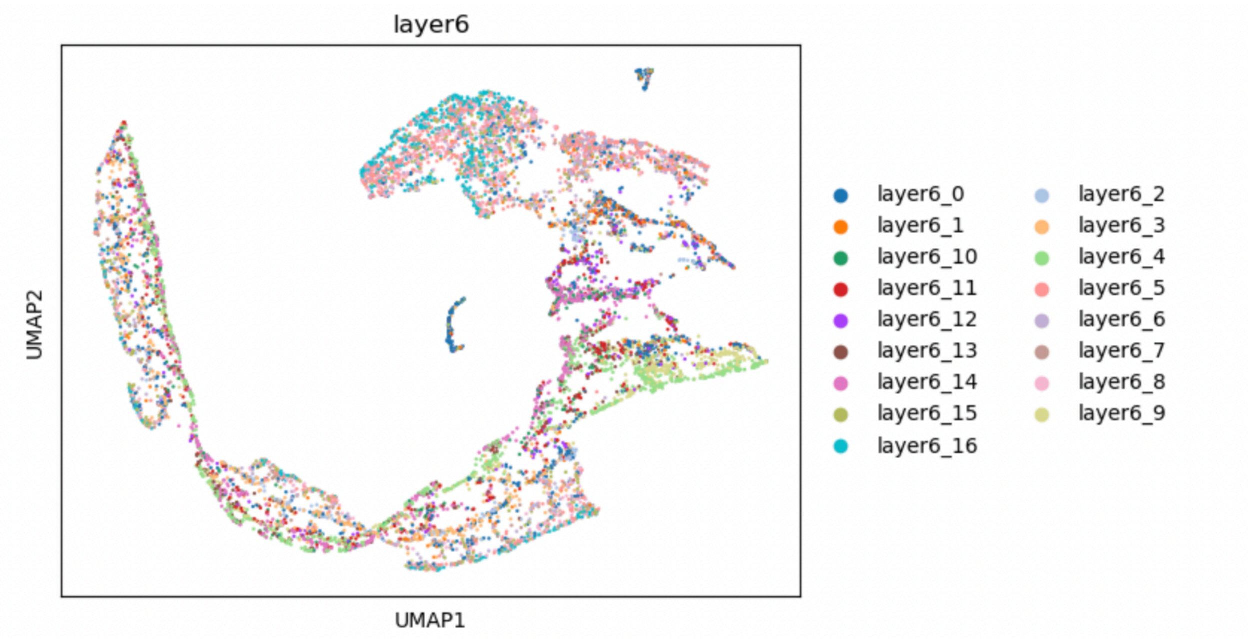

layer6
UMAP2
UMAP1
layer6_0
layer6_1
layer6_10
layer6_11
layer6_12
layer6_13
layer6_14
layer6_15
layer6_16
layer6_2
layer6_3
layer6_4
layer6_5
layer6_6
layer6_7
layer6_8
layer6_9


5

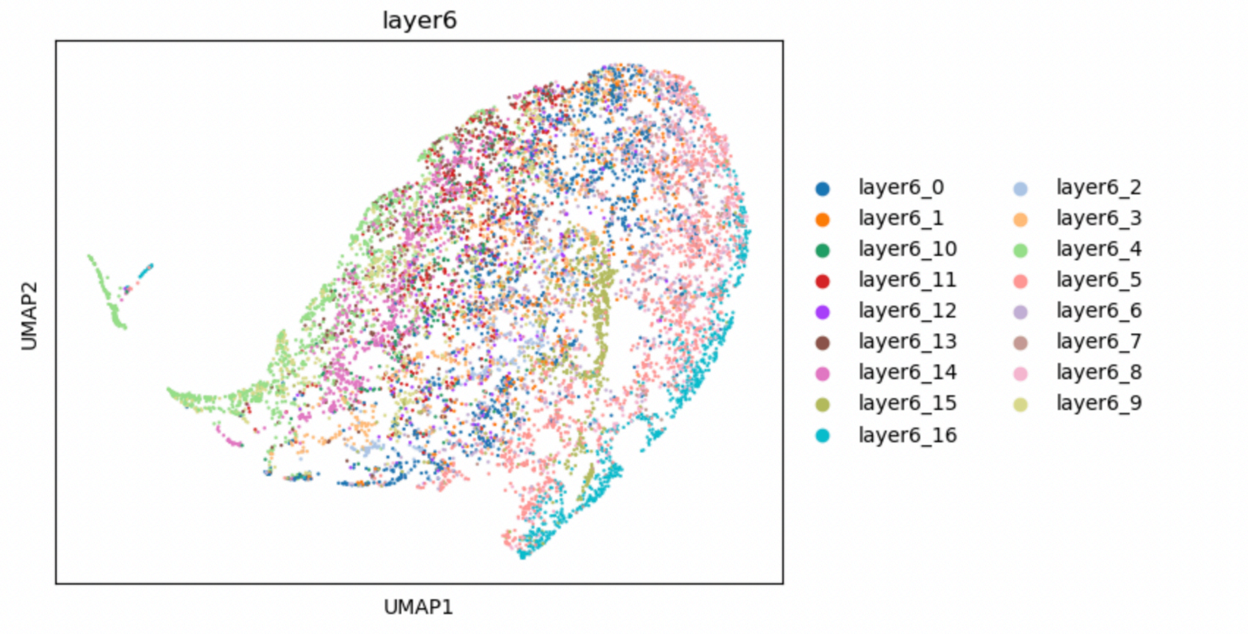

layer6
UMAP2
UMAP1
layer6_0
layer6_1
layer6_10
layer6_11
layer6_12
layer6_13
layer6_14
layer6_15
layer6_16
layer6_2
layer6_3
layer6_4
layer6_5
layer6_6
layer6_7
layer6_8
layer6_9


F
1

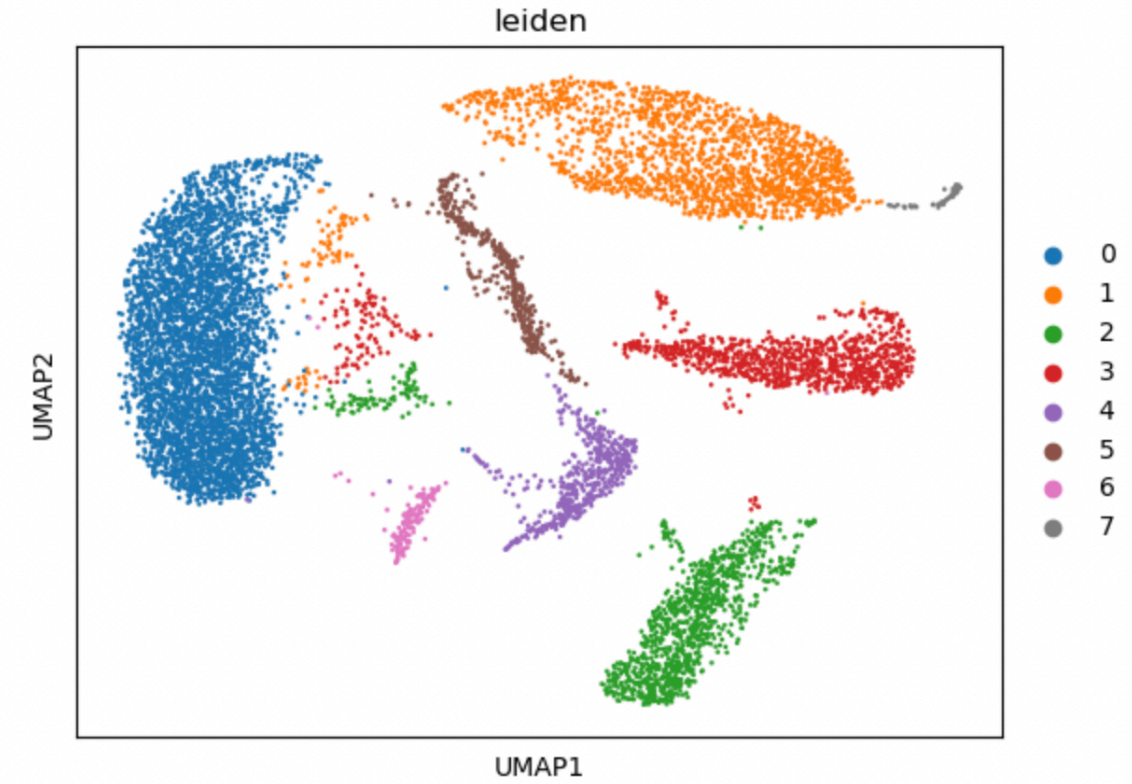
leiden
UMAP2
UMAP1
0
1
2
3
4
5
6
7

2

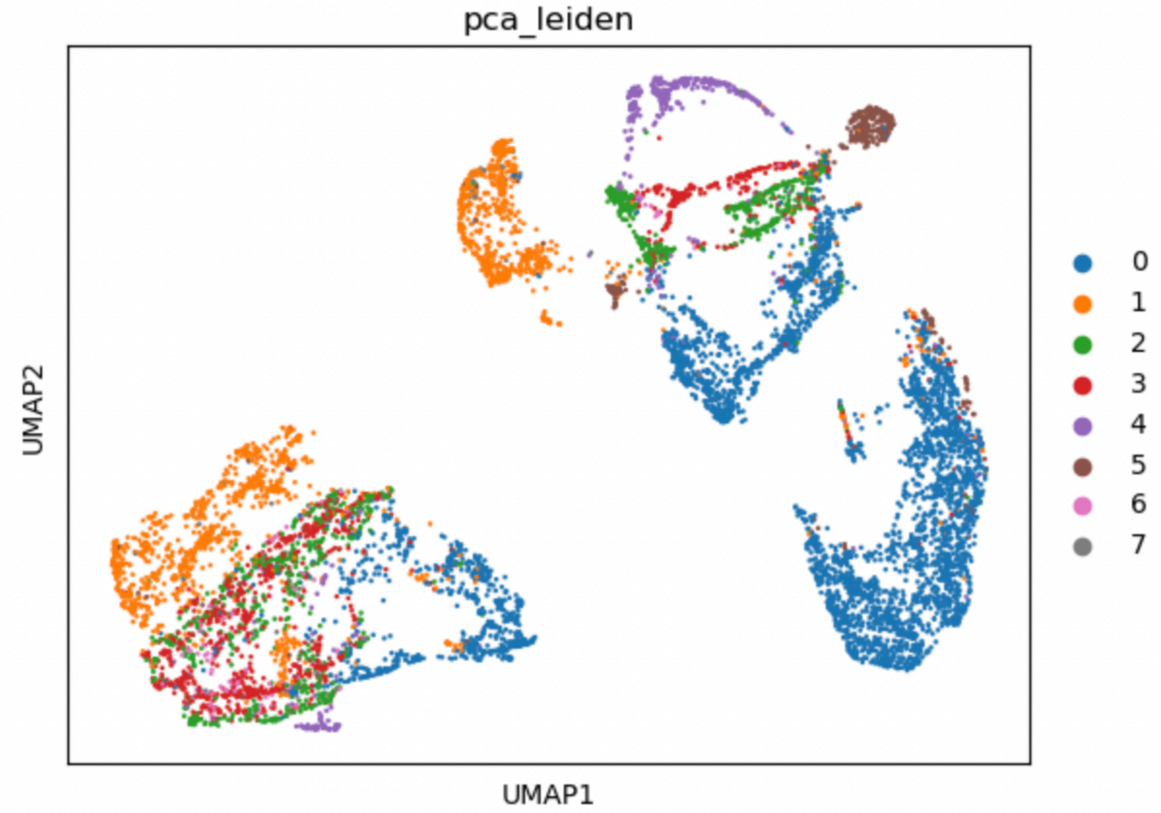
pca_leiden
UMAP2
UMAP1
0
1
2
3
4
5
6
7

3

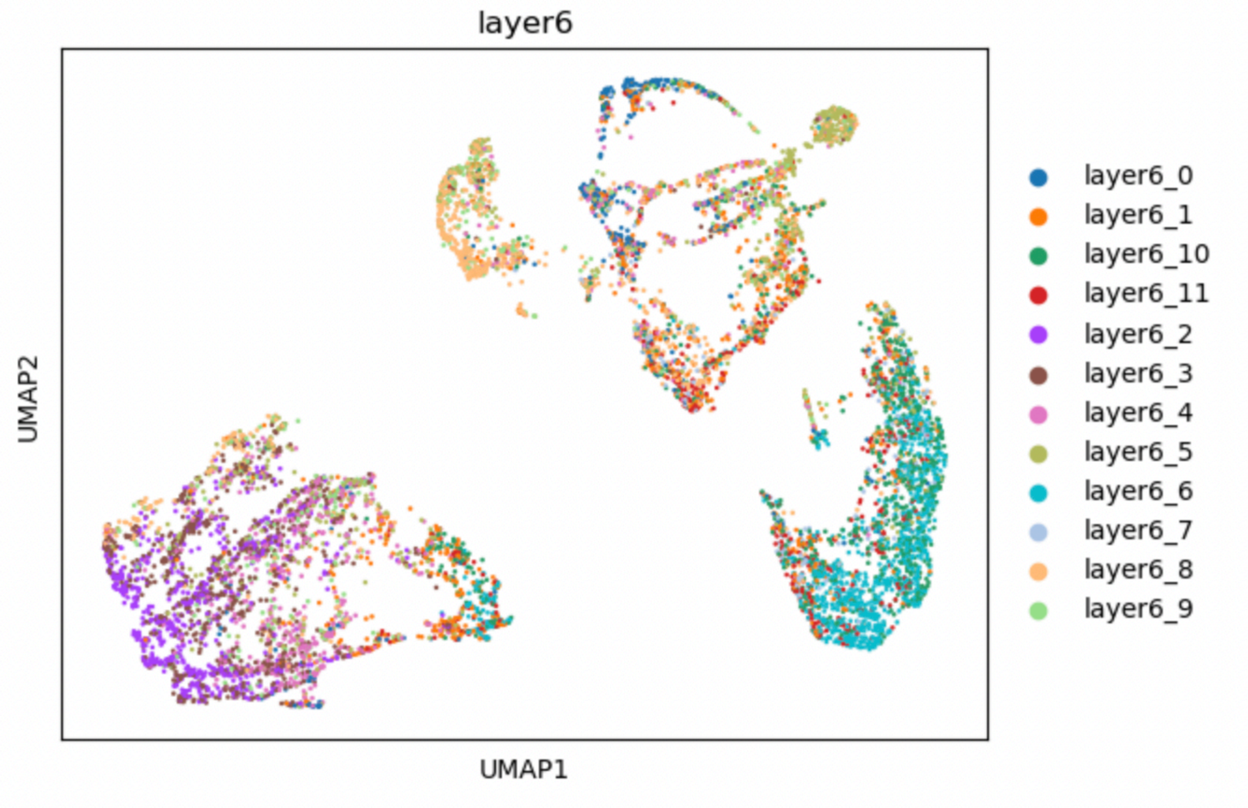
layer6
UMAP2
UMAP1
layer6_0
layer6_1
layer6_10
layer6_11
layer6_2
layer6_3
layer6_4
layer6_5
layer6_6
layer6_7
layer6_8
layer6_9

4

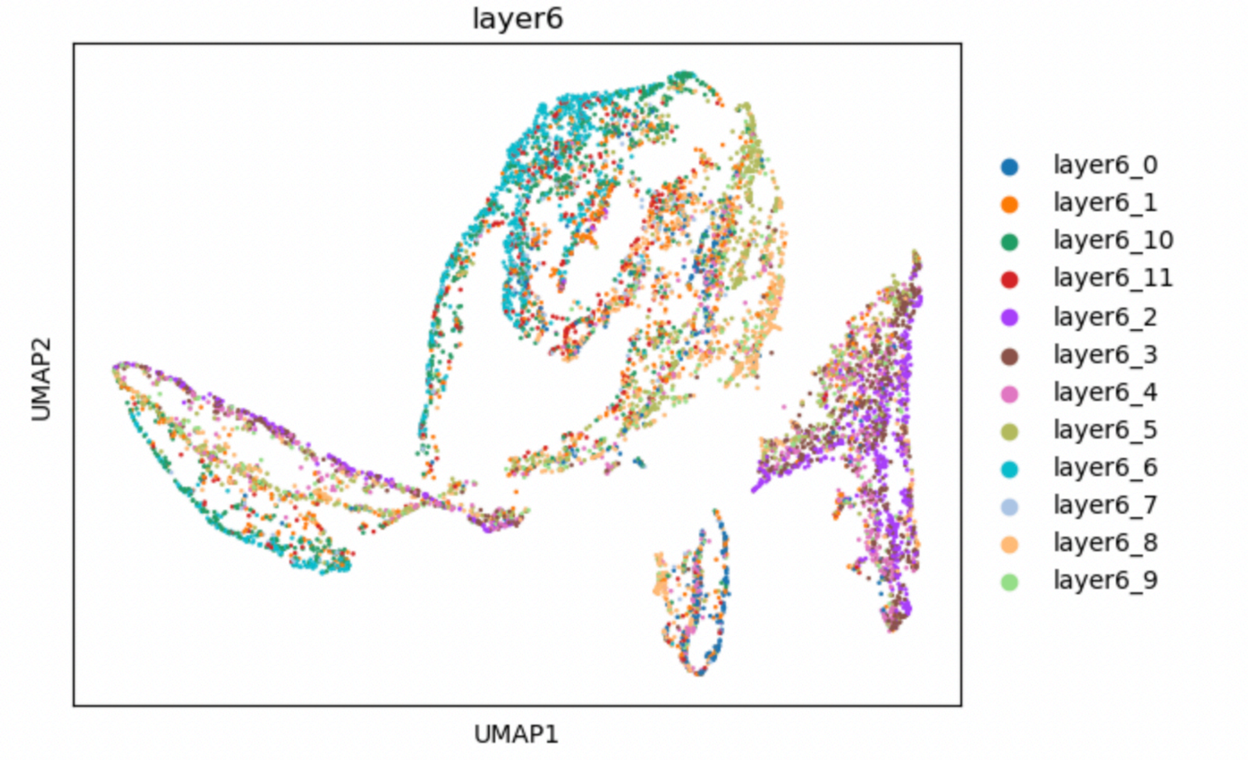
layer6
UMAP2
UMAP1
layer6_0
layer6_1
layer6_10
layer6_11
layer6_2
layer6_3
layer6_4
layer6_5
layer6_6
layer6_7
layer6_8
layer6_9

5

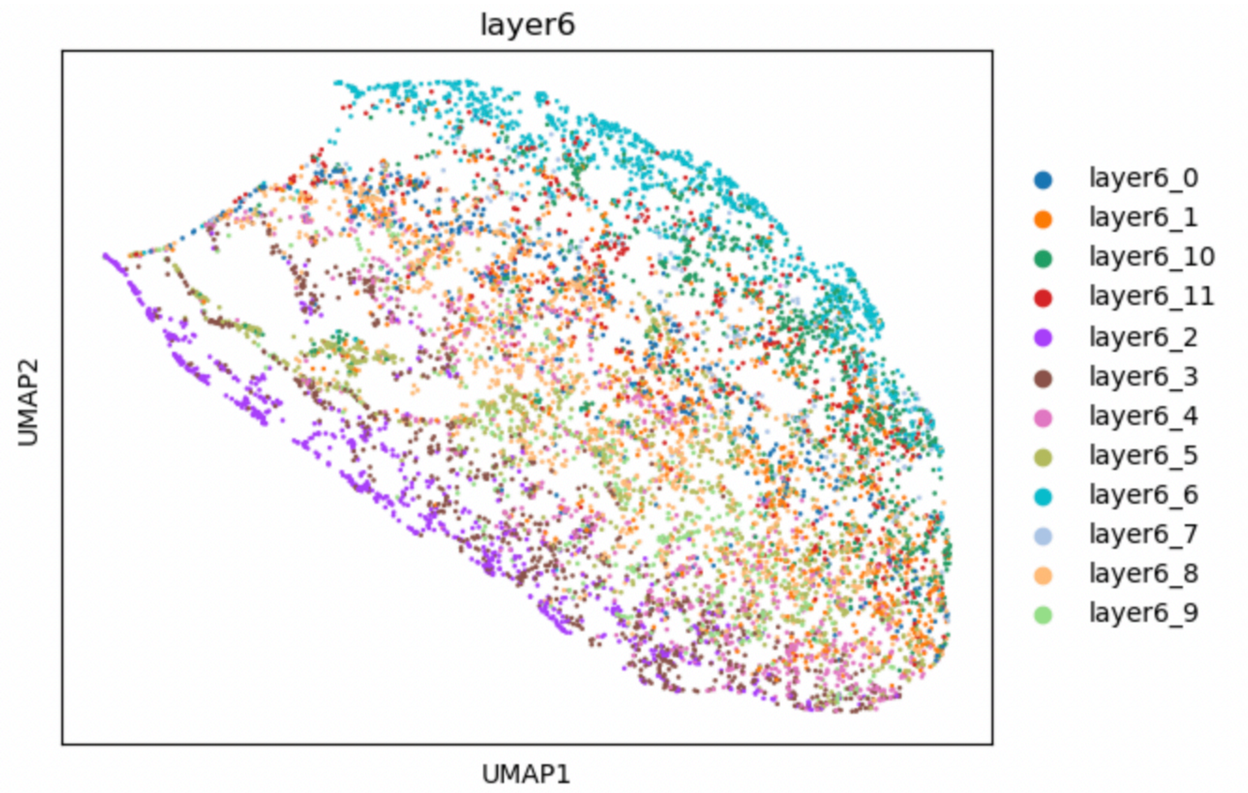


Legend: (1,2,3,4,5): PCA clusters and visualization, PCA projection on BW cluster graph, BW clusters on BW cluster graph, BW logged purged global trajectory graph, BW global trajectory graph by only purged matrix. (A, B, C, D, E, F): control1, control2, control3, AD1, AD2, AD3.
